# A first-in-class Wiskott-Aldrich syndrome protein (WASp) activator with anti-tumor activity in hematological cancers

**DOI:** 10.1101/2022.11.25.517686

**Authors:** Filippo Spriano, Giulio Sartori, Laura Barnabei, Alberto J. Arribas, Matilde Guala, Ana Maria Carrasco Del Amor, Meagan R. Tomasso, Chiara Tarantelli, Luciano Cascione, Gaetanina Golino, Maria E Riveiro, Roberta Bortolozzi, Antonio Lupia, Francesco Paduano, Samuel Huguet, Keyvan Rezai, Francesco Margheriti, Pedro Ventura, Greta Guarda, Giosuè Costa, Roberta Rocca, Andrea Cavalli, Giampietro Viola, Christoph Driessen, Emanuele Zucca, Anastasios Stathis, Beat Bornhauser, Stefano Alcaro, Francesco Trapasso, Susana Cristobal, Shae B. Padrick, Natalina Pazzi, Franco Cavalli, Francesco Bertoni, Eugenio Gaudio

**Affiliations:** Institute of Oncology Research, Faculty of Biomedical Sciences, USI, Bellinzona, Switzerland; SIB Swiss Institute of Bioinformatics, Lausanne, Switzerland; Chimete, Tortona, Italy; Department of Biomedical and Clinical Sciences, Cell Biology, Medical Faculty, Linköping University, Linköping, Sweden; Drexel University, College of Medicine, Department of Biochemistry and Molecular Biology, Philadelphia, PA, USA; Department of Pharmacy, University of Napoli Federico II, Napoli, Italy; Early Drug Development Group, Boulogne-Billancourt, France; Department of Woman’s and Child’s Health, University of Padova, Italy; Istituto di Ricerca Pediatrica IRP, Fondazione Città della Speranza, Padova, Italy; Net4science Srl, Magna Graecia University of Catanzaro, Catanzaro, Italy; University “Magna Græcia” of Catanzaro, Catanzaro, Italy; Tecnologica Research Institute and Marrelli Health, Biomedical Section, Stem Cells and Medical Genetics Units, Crotone, Italy; Institut Curie, Paris, France; Institute of Research in Biomedicine, Faculty of Biomedical Sciences, USI, Bellinzona, Switzerland; Kantosspital St Gallen, St Gallen, Switzerland; Oncology Institute of Southern Switzerland, Ente Ospedaliero Cantonale, Bellinzona, Switzerland; Children’s Hospital Zurich, Zurich, Switzerland; Ikerbasque, Basque Foundation for Sciences, Department of Department of Physiology, Faculty of Medicine and Nursing, University of the Basque Country, Spain

## Abstract

Hematological cancers are among the most common cancers in adults and in children. Despite significant improvements in therapies, many patients still succumb to the disease, therefore, novel therapies are needed. The Wiskott-Aldrich syndrome protein (WASp) family proteins regulate actin assembly in conjunction with the Arp2/3 complex, a ubiquitous nucleation factor. WASp is expressed exclusively in hematopoietic cells and exists in two allosteric conformations, auto-inhibited and active conformations. Here, we describe the development of EG-011, a first-in-class small molecule activator of the WASp auto-inhibited form. EG-011 possesses in vitro and in vivo anti-tumor activity as single agent in lymphoma, leukemia and multiple myeloma, including models of secondary resistance to PI3K, BTK and proteasome inhibitors. The in vitro activity was confirmed in a lymphoma xenograft. Actin polymerization induced by EG-011 was demonstrated with multiple techniques. Transcriptome analysis highlighted homology with drugs inducing actin polymerization.

**Key points:**

1. EG-011 is a novel small molecule with anti-tumor activity in hematological cancers, including resistant lymphoma and multiple myeloma models
2. EG-011 is a first-in-class small molecule activator of the auto-inhibited form of the Wiskott-Aldrich syndrome protein (WASp)

## Introduction

Hematological cancers are among the most common cancers in adults and in children ^1,2^. Improvements in patients outcome are due to the introduction of therapies against novel targets (for example inhibitors of BTK, PI3K, BCL2 of EZH2) or drugs with novel mechanism of action on already known targets (for example, anti-CD19 chimeric antigen receptor T-cells, antibody drug conjugates, or bispecific antibodies) ^3^. Despite access to these therapies and the improvements in the management of patients, still too many individuals succumb due to the emergence of relapses frequently refractory to standard therapies ^1,4,5^.

The human Wiskott-Aldrich Syndrome (WAS) gene is almost exclusively expressed in cells belonging to the hematopoietic lineage ^6^. Disruption of its protein (WASp) through loss-of-function (LOF) or activating gain-of-function (GOF) mutations give rise to Wiskott-Aldrich syndrome (WAS) and in X-linked neutropenia (XLN), respectively ^7–11^. In XLN, hyperactivated WASp causes increased actin polymerization, causing genomic instability and apoptosis in lymphocytes ^12^. In WAS patients, depletion of WASp protein or activity disrupts actin polymerization in T cells ^13^. WASp directly interacts with FYN ^14^ and BTK ^15^, which are proteins involved in B cell receptor and Toll-like receptor signaling^3^. B cells without functioning WASp have defective actin cytoskeletons with reduced motility and aggregates compared to normal B cells ^16^. Deletion of WASp in B cells also leads to production of autoantibodies and the development of autoimmunity, as in Wiskott-Aldrich syndrome patients ^17^, and *WAS* mutations are associated with the development of lymphomas in around 15% of cases^18^. The complexity of WASp regulation and functions is underlined by data in ALK-positive anaplastic large cell lymphoma (ALCL), reporting both an oncogenic ^19^ and a tumor suppressive role for the protein^20^.

These identify the protein both as an oncogene and a tumor suppressor ^20^.

WASp belongs to a family of proteins that share the VCA (verprolin homology, central, acidic) motif in their C-terminal portion ^21^. WASp and the other members of the family (N-WASP, WAVE/SCAR1,2 and 3, WASH, JMY and WHAMM) are involved in the actin assembly and cytoskeleton reorganization, using their VCA domains to promote nucleation of actin filaments by the Arp2/3 complex ^21^. The activity of WASp is regulated through a well-studied allosteric mechanism, wherein WASp exists in two allosteric conformations. In the first (autoinhibited) conformation, an N-terminal domain binds to and sequesters the C-terminal VCA domain, in a fashion coupled to folding of the protein^22–24^. The autoinhibited form is then activated by the competitive binding of the small GTPase Cdc42 ^25^, binding of the phospholipid PtdIns(4,5)P2 ^26^ or phosphorylation ^27^. Additional layers of regulation occur through clustering of WASp molecules into high potency complexes^28,29^.

Small molecules affecting WASp functions have been reported. The small molecule wiskostatin inhibits WASp by inducing folding of the GBD into its autoinhibited conformation stabilizing WASp into its autoinhibited conformation^30^. Another reported compound (SMC#13) promotes WASp degradation and has anti-tumor activity in lymphoma and leukemia models ^31^. Thus, small molecules may mimic the effects of WASp mutants and selectively lead to cell death in hematological lineages, indicating WASp as a new target not yet explored in the clinical setting.

Here, we describe the fortuitous identification of a first-in-class WASp activator (EG-011) with specific activity in various hematological malignancies.

## Methods

### Cell lines

Sixty-two cell lines derived from human and non-human lymphomas (Supplementary Table 1) were cultured according to the recommended conditions. All media were supplemented with fetal bovine serum (10/20%), Penicillin-Streptomycin-Neomycin (~5,000 units penicillin, 5 mg streptomycin and 10 mg neomycin/mL, Sigma Aldrich, US) and L-glutamine (1%). Cell line identity was authenticated by short tandem repeat DNA profiling ^32^. All the experiments were performed within one/two month from being thawed. Cell lines were periodically tested to confirm Mycoplasma negativity using the MycoAlert mycoplasma Detection Kit (Lonza, Visp, Switzerland). Detailed methods for additional cell lines derived from solid tumors and leukemia are provided in Supplementary Materials and Methods.

### Compounds

EG-011 was synthetized by Chimete (Tortona, Italy) (See Results section and supplementary results section). Rituximab (Roche, Switzerland) was dissolved in physiological solution at the concentration of 10 mg/ml. Additional anti-cancer agents were purchased from Selleckchem (TX, USA) and prepared in DMSO at the stock concentration of 100 mM.

### MTT proliferation assay, Cell cycle and Apoptosis assay

Full methods are provided in Supplementary Materials and Methods.

### Patient-derived xenografts

Drug responses in ALL patient-derived xenografts (PDX) were analyzed as previously described ^33^. Full methods are provided in Supplementary Materials and Methods.

### Drug combinations

OCI-LY-1, OCI-LY-8, REC1, MINO and TMD8 cell lines were exposed (72 h) to increasing doses of EG-011 alone or in combination with increasing doses of FDA approve compounds (Rituximab, bendamustine, Ibrutinib, lenalidomide and ABT-199), followed by MTT assay. Ibrutinib and lenalidomide were tested in MCL and ABC-DLBCL (TMD8) were they showed clinical response. Synergism was calculated using the Chou-Talalay combination index (CI) and the MuSyC algorithm ^34,35^. CI values are referred as median CIs for all the different combinations; additive effect (CI 0.9-1.1), synergism (CI 0.3-0.9), strong synergism (CI<0.3) and antagonism/no benefit (CI > 1.1). Efficacy values are calculated for every different concentration combination of the two drugs. Potency is calculated on IC_50_s and one value for each curve is obtained. Concentrations that gave 10% or less of proliferation already with the single agent were discarded for further analyses.

### Xenografts experiments

NOD-Scid (NOD.CB17-*Prkdcscid*/NCrHsd) mice maintenance and animal experiments were performed under institutional guidelines established for the Animal Facility and with study protocols approved by the local Cantonal Veterinary Authority (No. TI-20-2015). Full methods are in Supplementary Materials and Methods).

### Primary healthy PBMCs

Peripheral blood was obtained from two healthy donors. Peripheral blood mononuclear cells (PBMCs) were isolated by Ficoll density-gradient centrifugation and then cultured in the presence of DMSO or EG-011 (two doses: 1 and 10 μM). After 24 and 48 hours, the percentage of apoptotic cells was determined by staining of the cells with Annexin V–FITC

### Gene expression profiling

REC1 cell line was exposed to DMSO and to EG-011(500 nM) for 8 hours. Gene Expression Profiling (GEP) was done using the HumanHT-12 v4 Expression BeadChip (Illumina, San Diego, CA, USA). Data processing and statistical analysis was performed using R/Bioconductor ^36^, as previously described ^37^. Transcript mapping was based on HG19 using manufacturer’ supplied annotation. Data were analyzed using the omics playground platform (https://bigomics.ch)

### Kinase assay screenings

EG-011 was screened against 468 human kinases at the concentration of 1000 nM by using the KINOMEscan assay, ATP independent, provided by Discoverx (CA, USA). EG-011 was also tested at the concentration of 100 and 1000 nM against 320 kinases by running a regular ATP related assay, Wildtype-Profiler, provided by ProQinase (Freiburg, Germany).

### Thermal proteome profiling (TPP)

Experiments were performed as described in Franken et al. ^38^ with some modifications (see Supplementary Materials and Methods).

### Functional WASp experiments

Actin was purified from rabbit muscle according to the Spudich method^39,40^. Briefly, an extract of rabbit muscle acetone powder (Pel-Freeze Biologicals) in Buffer G (2 mM Tris-HCl pH 8, 200 μM ATP, 0.5 mM DTT, 0.1 mM CaCl2, 1 mM sodium azide) was polymerized by the addition of magnesium chloride and potassium chloride. Filaments were harvest by centrifugation, depolymerized by dialysis into Buffer G and then purified by size exclusion chromatography using Superdex 200pg in Buffer G. Pyrene actin was prepared by labeling polymerized actin with a ten-fold molar excess of N-(1-pyrene) iodoacetamide for 16 hours at 4°C. Labeled filaments were then collected, depolymerized and purified by size-exclusion chromatography. Actins were stored in Buffer G in continuous dialysis at 4°C.

Active and autoinhibited WASp were purified as described ^41,42^. Briefly, pET15 plasmids encoding for WASp GBD-P-VCA (active) and WASp BGBD-VCA (autoinhibited)^41^ were expressed as His6 tagged fusions in BL21(DE3) T1^R^ E. coli, inducing overnight at 20°C with 1 mM IPTG. Cells were lysed by extrusion (Emulsiflex C5, Avestin Inc., Ottawa, Canada) and centrifuged at 17,000 RPM in a JA-20 rotor for 45 minutes. The supernatant was applied to Nickel Sepharose FF beads, washed and eluted. The Ni column eluate was applied to a SOURCE15Q anion exchange chromatography column, and eluted with a 0-50% Buffer B gradient over 30 CV (Buffer A= 20 mM Imidazole pH 7, 1 mM DTT; Buffer B= 20 mM Imidazole pH 7, 1 mM DTT, 1 M Sodium Chloride). Appropriate fractions were determined by SDS-PAGE analysis and applied to a HiLoad 26/600 Superdex 75pg column (Cytiva) developed in a buffer of 150 mM potassium chloride, 2 mM MgCl_2_, 1 mM EDTA, 10 mM Imidazole pH 7. Fractions were pooled, supplemented with glycerol to 20% w/v, aliquoted, flash frozen in liquid nitrogen and stored at −80°C. Arp2/3 complex was purified as previously described ^43^.

### Pyrene actin assembly assays

Pyrene actin polymerization assays were performed with 2 μM total actin (10% pyrene labelled) using established methods^39^. An actin mix was prepared by mixing unlabeled actin monomers with pyrene labelled actin monomers. 10E1M buffer (10 mM EGTA pH 8, 1 mM MgCl_2_) was added to 1/9^th^ volume and rest of volume was filled by buffer G-Mg (2 mM Tris-HCl pH 8, 200 μM ATP, 0.5 mM DTT, 0.1 mM MgCl_2_) to make solution A. Solution B contains 1/9^th^ volume 10x KMEI (500 mM potassium chloride, 20 mM MgCl_2_, 10 mM EDTA, 100 mM Imidazole pH 7) and enough Arp2/3 complex, WASp and/or EG-011 to yield the indicated concentrations after dilution with solution A. EG-011 stock was stored in DMSO at 10 mM and was diluted to a working stock of 50 μM in 1x KMEI immediately before use. Solutions A and B were mixed at a 1:1 ratio in a well of a 96 well half-area plate. Measurement of pyrene fluorescence began with an average deadtime of around 60 seconds. Fluorescence intensity was recorded every 2.5 seconds, exciting at 365 nm, 5 nm bandpass and detecting emission at 407 nm with 5 nm bandpass, with a 410 nm dichroic mirror separating the excitation and emission light paths.

### Cellular PK of EG-011

Intra and extracellular concentrations of EG-011 analysis were done as follow. Cells were seeded at 1×10^6^ cells/mL and exposed to 96.5 ng/mL (0.2 μM) EG-011 for 0, 5, 15, 30, 60, 120 and 360 min. At each time point, cells were collected and centrifuged at 2000 rpm for 5 minutes at 4C. Supernatant (5 mL) was collected, and the cell pellet was washed with PBS and stored at −20C. EG-011 extracellular and intracellular concentrations were analyzed in cell supernatants and pellets respectively, using Ultra Performance Liquid Chromatography with tandem mass spectrometry.

### Immunofluorescence

For immunofluorescence methods, we used VL51 sensitive cell line since REC1 cell line was not compatible with the attaching protocol used. Full methods are in Supplementary Materials and Methods.

## Results

### EG-011 is a novel small molecule

We modified the central scaffold of the FDA approved BTK inhibitor ibrutinib (Figure 1A, B), creating a new molecule, termed EG-011, containing an aromatic heterocyclic nucleus pyrido [3,2-d] pyrimidine-2,4 (1H, 3H) -dione instead of the ibrutinib aromatic heterocyclic nucleus 1H-pyrazolo [3,4-d] pyrimidin-4-amino. EG-011 differed with the 4phenoxyphenyl substituent in position 3 replaced with the 4phenoxybenzyl in position 1 and the piperidine substituent in position 1 shifted to position 3 (Figure 1B). The synthetic route to EG-011 was distinct from ibrutinib (Figure 1C) and is reported in the supplementary results section.

**Figure 1.**
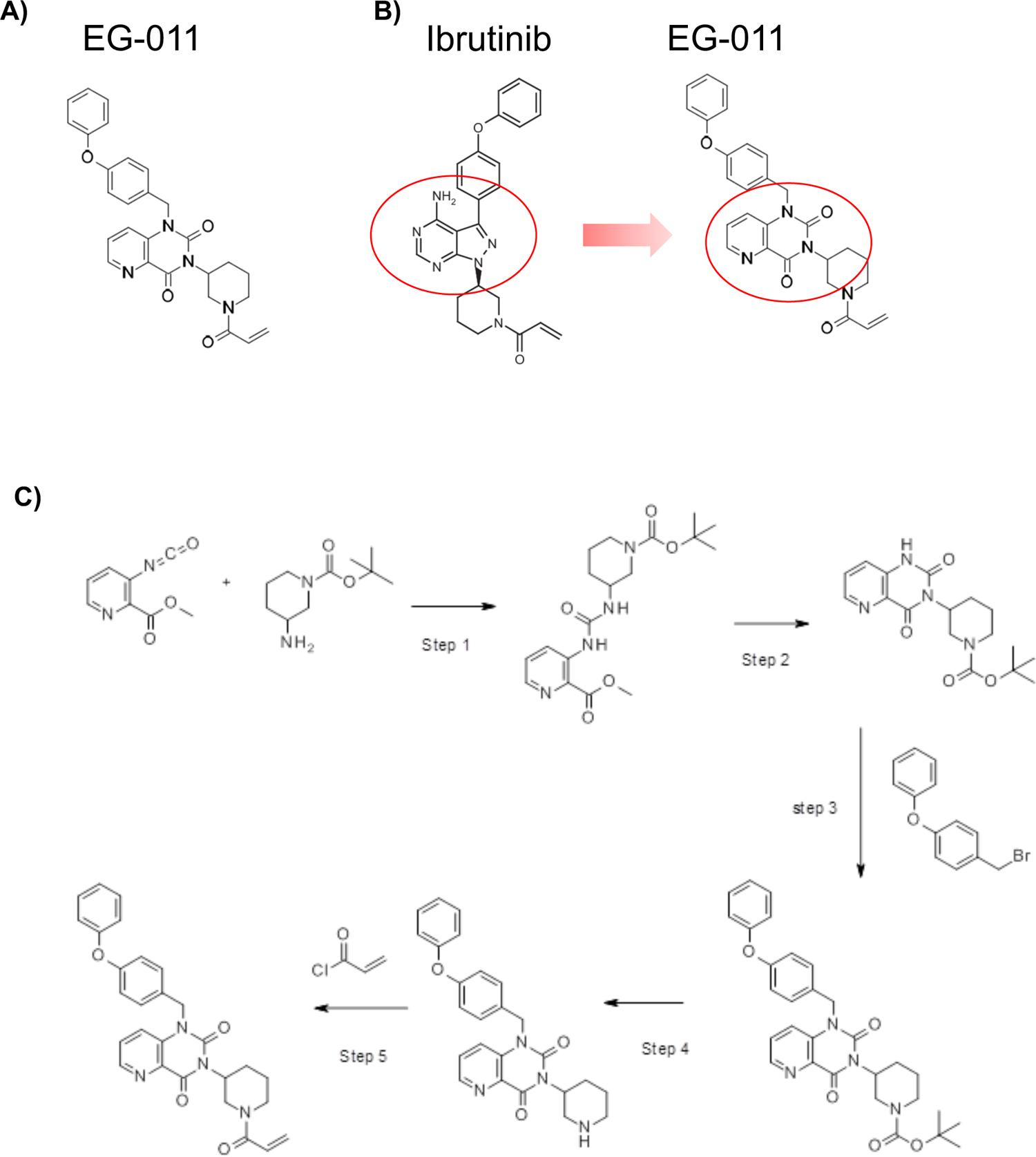
EG-011 chemical structure and synthetic route. **A**, EG-011 chemical structure; **B**, Structural modifications between ibrutinib and EG-011. **C**, Synthetic route of EG-011.

### EG-011 has anti-tumor activity only among hematological cancers

We tested the potential anti-lymphoma activity of EG-011 in a panel of 62 lymphoma cell lines, derived from various histological subtypes. EG-011 showed anti-lymphoma activity with a median IC_50_ of 2.25 μM (95% C.I. 1-5 μM) (Figure 2A,B; Supplementary Table 1). A higher activity was observed in a group of 21 cell lines that had a median IC_50_ of 250 nM (95% C.I. 40-600 nM). Among these there were 11 germinal center B cell (GCB) diffuse large B cell lymphomas (DLBCL) (sensitive n=11/21, resistant n=9/41, p < 0.05), four mantle cell lymphoma (MCL) (sensitive n=4/21, resistant n=6/41, p n.s.) and three marginal zone lymphoma (MZL) (sensitive n=3/21, resistant n=2/41, p n.s.).

**Figure 2.**
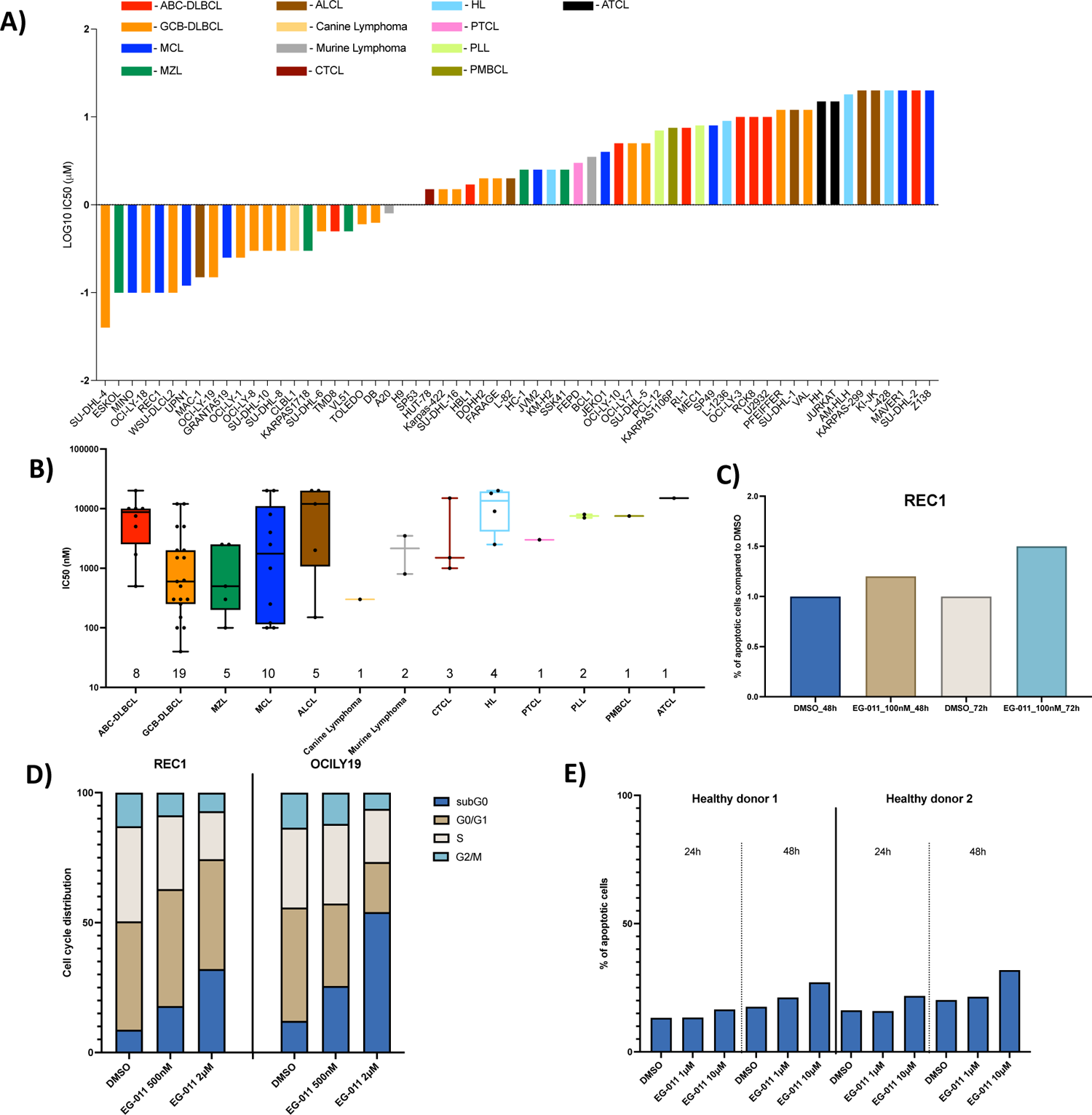
EG-011 has strong in vitro anti-lymphoma activity. **A**, *In vitro* activity, represented as IC_50_s calculated after 72 h of treatment, in 62 lymphoma cell lines. Cell lines are colored differently based on the different histological subtype. **B**, IC_50_s distribution after EG-011 treatment among the different subtypes of lymphoma. **C**, Apoptosis induction after 48 or 72 h EG-011 treatment (100 nM). **D**, Cell cycle changes after 72 h EG-011 treatment (500 nM and 2 μM) in two sensitive lymphoma cell lines. **E**, Apoptosis induction by EG-011 treatment at 24 and 48 h in two primary cells from healthy control patients.

No activity was observed in 11 acute leukemias cell lines (Supplementary Figure 1) but 7 out of 12 primary cells derived from acute lymphoblastic leukemia (ALL) were sensitive to EG-011 with IC_50_ values between 0.3-4.6 μM after 72 h of exposure. The remaining five displayed IC_50_ higher than 20 μM (Supplementary Figure 2).

EG-011 did not show any anti-proliferative activity in a panel of 23 solid tumor cell lines (IC_50_ > 10 μM), with only one cell line (head and neck tumor) sensitive with IC_50_ of 3.5 μM (Supplementary Figure 2).

Comparing the activity of EG-011 in lymphoma cell lines and solid tumors, it is clear that EG-011 has activity specifically among hematological cancers (Supplementary Figure 1).

The observed anti-proliferative activity of EG-011 is characterized by induction of cell death, rather than to cell cycle arrest, as indicated by surface exposed Annexin V and the dose-dependent increase in sub-G0 component (20-55%) observed in two sensitive lymphoma cell lines (OCI-LY-19 and REC1) exposed to the compound (500 nM and 2 μM; 72 h) (Figure 2C, D). No cytotoxicity was seen in PBMCs from two healthy donors after treatment with EG-011 at 1 and 10 μM for 24 h and 48 h (Figure 2E).

### EG-011 is active in cell lines resistant to FDA approved compounds

We tested EG-011 in models of secondary resistance to FDA approved PI3K and BTK inhibitors developed in our laboratory from splenic MZL cell lines ^44–46^. The anti-tumor activity of EG-011 was maintained or even increased in the resistant cell lines, both in terms of IC_50_ and area under the curve (AUC). EG-011 was especially active in VL51 idelalisib resistant cell lines compared to their parental counterparts (IC_50_ 100 nM / AUC 895.7 vs IC_50_ 500 nM / AUC 1,124). (Figure 3A, B, Supplementary Figure 3 A). To further asses the lack of cross resistance with others anti-cancer agents, we tested EG-011 in various multiple myeloma (MM) models with acquired resistance to proteosome inhibitors ^47–49^ (Figure 3C, D, E, F, Supplementary Figure 3 B). The anti-tumor activity of EG-011 was increased in MM resistant cells. The AMO-1 carfilzomib resistant cell line showed an IC_50_ to EG-011 treatment that was more than 20 times lower than the carfilzomib sensitive parental cell line in terms of IC_50_ (IC_50_ 250 nM / AUC 217,115 vs IC_50_ 6 μM / AUC 624,857). The L363 bortezomib resistant line showed an IC_50_ around ten times lower than the sensitive parental cell lines (IC_50_ 1 μM / AUC 357,564 vs IC_50_ 12 μM / AUC 852464). A 4-fold increase in sensitivity was observed in both RPMI-8266 bortezomib resistant (IC_50_ 2.5 μM / AUC 223,444) and in RPMI-8266 carfilzomib resistant cells (IC_50_ 2.5 μM / AUC 430,533) compared to parental cell lines (IC_50_ 10μM /AUC 591,981).

**Figure 3.**
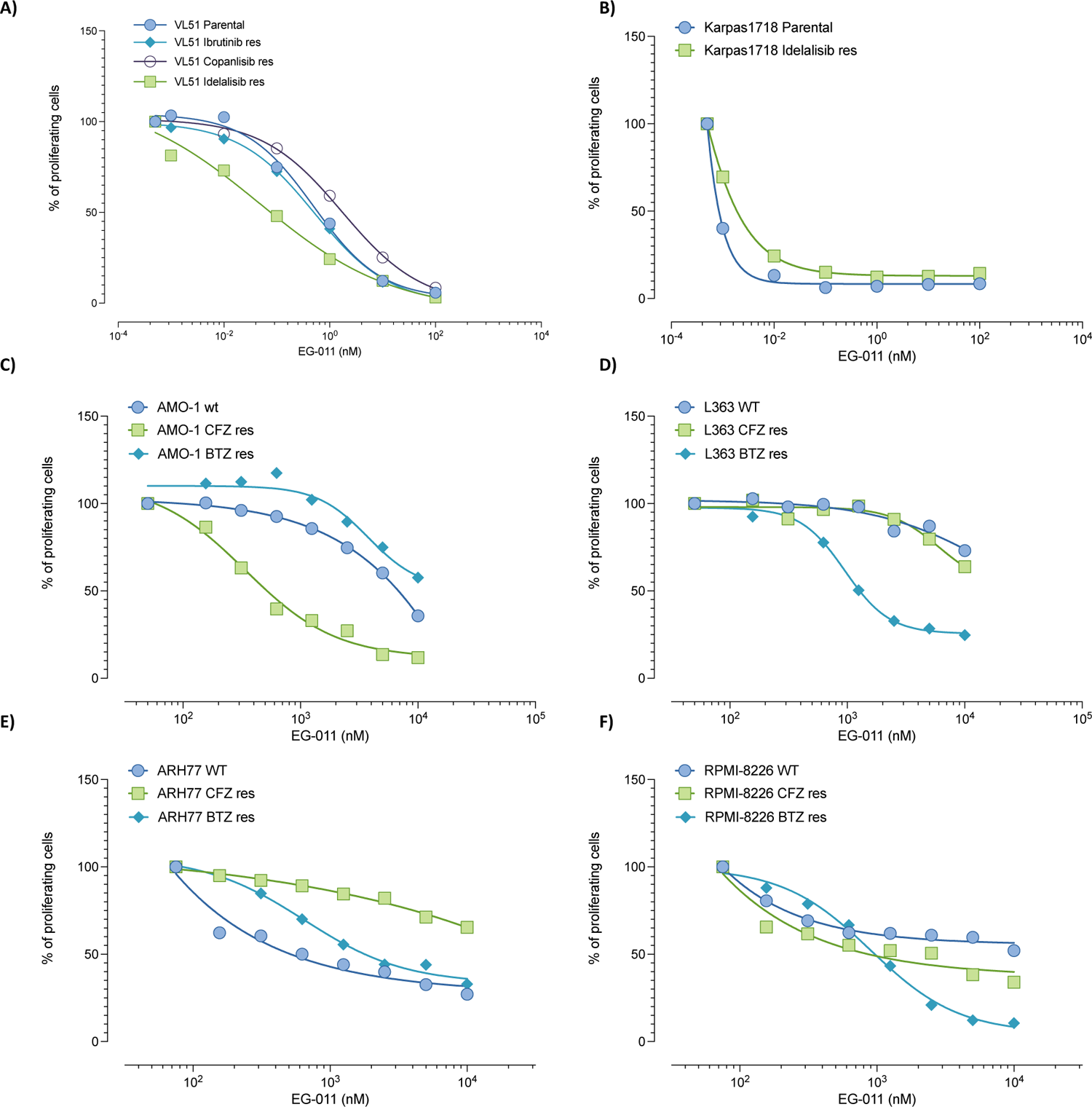
EG-011 is active in in models of secondary resistance to FDA approved compounds derived from splenic marginal zone lymphoma (A-B) and from multiple myeloma (C-F). Dose response curve after 72 h EG-011 treatment in cell lines with acquired resistance to FDA approved compounds compared to parental cell lines. Splenic marginal zone lymphoma cell lines: **A**, VL51, parental, resistant to idelalisib, ibrutinib or copanlisib; **B**, Karpas1718 parental or resistant to idelalisib. Multiple myeloma cell lines: **C**, AMO-1; **D**, L363; **E**, ARH77; **F**, RPMI-8226 parental, resistant to proteosome inhibitors carfilzomib (CFZ) and bortezomib (BTZ).

These data indicate that EG-011 was active also after acquired resistance to different FDA approved agents.

### EG-011 has *in vivo* anti-lymphoma activity

The observed *in vitro* anti-tumor activity of EG-011 was confirmed *in vivo* using the REC1 MCL cell line in a mouse xenograft model. EG-011 was administered at 200 mg/kg once per day, 5 days per week and compared to vehicle control. EG-011 delayed tumor growth (volume) versus control (Day 6, Day 7, Day 9, p < 0.05) and final tumor weight (Figure 4A, B). EG-011-treated tumors were 2.2-fold smaller than controls (p < 0.001). Treatments were well tolerated in mice, without significant signs of toxicity. Throughout treatment, mice were well-conditioned with a body condition score ^50^ BC3 for all groups.

**Figure 4.**
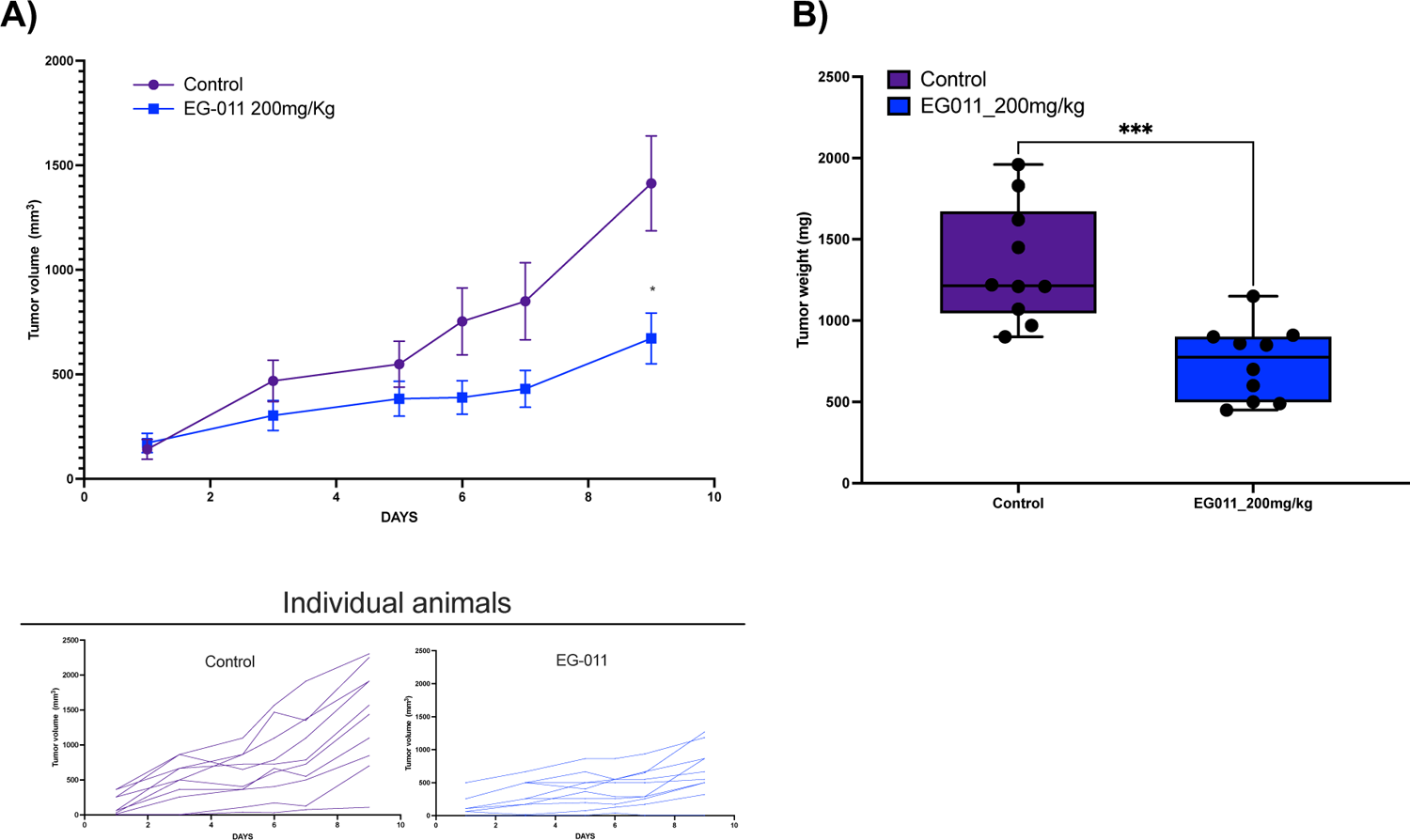
EG-011 has *in vivo* anti-lymphoma activity. **A,** *in vivo* activity of EG-011 (200 mg/Kg, IP, no.= 10 mice) compared to control vehicle (IP, no.= 9 mice) in NOD-SCID mice. Upper graph represents the mean tumor volume (mm^3^) with standard error mean (SEM) for each day. Lower graph, represents the growth of each individual animals under control vehicle or EG-011 treatment. **B**, Tumor weights (mg) of control and EG-011 treated mice at the end of the treatment (day 9). P-values are calculated with Mann Whitney test. * p-value < 0.05; ***, p-value <0.0001.

### Differences in cellular uptake do not explain the lack of activity in solid tumors

We analyzed the cellular PK properties of EG-011 in sensitive and resistant cell lines to address whether the lack of activity of EG-011 in solid tumor models could be sustained by differences in compound’s intake and kinetics. The colorectal cancer cell line HCT-116 was used as a resistant model (IC_50_ > 10 μM) and the MCL cell line REC1 as a sensitive model. Cellular uptake of EG-011 was rapid (< 5 min) in both sensitive and resistant cell lines, with a mean concentration of 10-20 ng/ml every 10^6^ cells (20-40 nM) after 6h exposure (Supplementary Figure 4). Extracellular levels of EG-011 were stable from treatment initiation for up to 6h exposure (~50 ng/mL in both cell lines. We observed a ratio of ~5 between intracellular and extracellular concentrations in both cell lines. In addition, we did not observe an active drug ejection of EG-011 in HCT-116 cells (Supplementary Figure 4). These data indicated that the lack of activity in solid tumors was due to neither a different cellular uptake nor a higher ejection of the compound.

### Transcriptome changes due to EG-011

We exposed a sensitive cell line (REC1) to EG-011 and looked at the transcriptome changes after 8h (Supplementary Table 2). We identified, among others, a downregulation in MYC targets, mitotic spindle assembly genes involved in actin filaments organization. Comparing the genes modulated by EG-011 with publicly available data of other drugs (L1000^51^ and GDSC^52^), the new small molecules behaved similarly to microtubules-stabilizing agents and HDAC inhibitors. Among the top correlated drugs there were indeed docetaxel and epothilone B (Supplementary Figure 5; Supplementary Table 3). Lymphoma and pro-survival genes such as CXCR5, NFKBID, and BCL2A1 were among the top downregulated genes.

### EG-011 is the first in class WASp activator

As above mentioned, EG-011 was designed by modifying the BTK inhibitor ibrutinib, however its pattern of activity did not correlate with ibrutinib’s activity (Supplementary Figure 6), indicating that EG-011 did not act via inhibiting the kinase. To assess whether additional kinases were inhibited by the compound, we performed two screenings. First, EG-011 (1 μM) was compared to DMSO with a competition binding assay against 450 human kinases and disease relevant mutants (KINOMEscan). Second, EG-011 (100 nM, 1 μM) was compared to DMSO with a radiometric protein kinase assay against 320 human kinases (PanQinase Activity Assay). No effect against any kinase was observed (Supplementary Table 4) indicating that EG-011 was not a kinase inhibitor.

To identify possible EG-011 targets in an unbiased manner, we then applied the thermal proteome profiling (TPP) technique ^38,53^. The latter relies on the principle that, when subjected to heat, proteins denature and become insoluble, while upon interactions with small molecules proteins can change their thermal stability. TPP was used by applying label-free quantitative mass spectrometry, facilitating the analysis of the changes of the melting profile within a complex mixture or across an entire proteome in a single experiment. TPP was performed on proteins extracted from EG-011-treated REC-1 cells (10 μM) and the corresponding control with a vehicle. The analysis of the EG-011 protein interactions identified 48 potential protein targets, 8 stabilized and 40 destabilized (Supplementary Figure 7, Supplementary Table 5). WASp was the most highly destabilized protein by EG-011 (Supplementary Table 5, Figure 5A).

**Figure 5.**
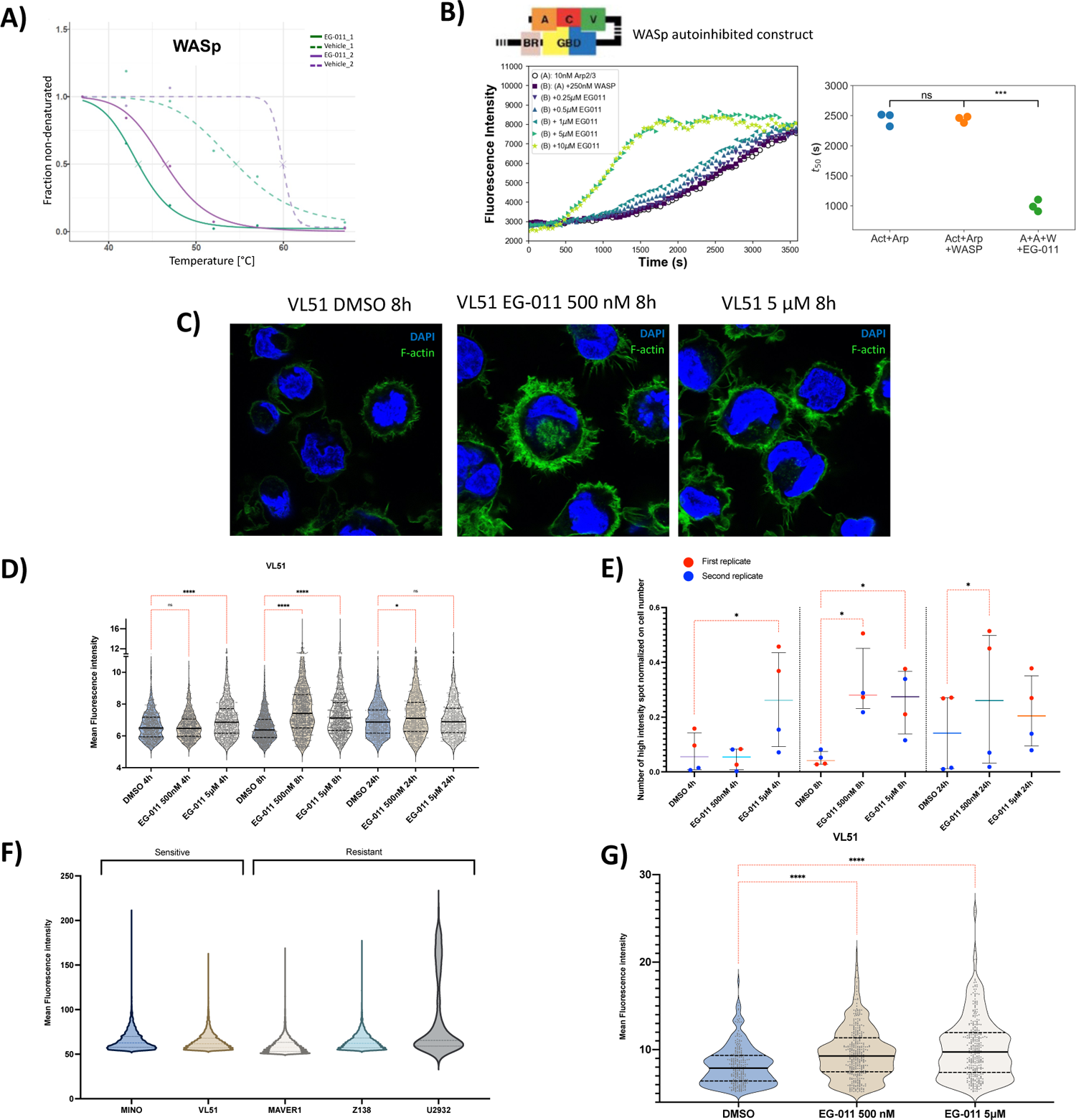
EG-011 is the first in class WASp activator. **A,** Melting curves of WASp under DMSO (dashed lines) or EG-011 (solid lines) treatment. Protein lysate from an EG-011 sensitive cell line (REC1) was subjected to the thermal shift assay performed at temperatures between 37 °C to 67 °C in presence or absence (vehicle) of EG-011. The unfolding profile shown a high decrease of solubility of WASp protein in presence of EG-011 vs. vehicle. Melting temperature is shown with an “X” for each melting curve. Two biological replicates. **B**, Pyrene actin polymerization measured as fluorescence intensity, in the presence of the auto-inhibited construct of WASp. Actin polymerization in the absence or presence of EG-011 at five different concentrations (10, 5, 2.5, 1, 0.5, 0.25 μM). Tukey-HSD comparisons. **C**, Representative confocal images of sensitive lymphoma cell line treated for 8h with DMSO or EG-011 (500 nM and 5 μM). Quantification of mean fluorescence intensity (ImageJ) per cell line in a 20X image (Leica widefield microscope) with n of around 200 cells. Experiments were performed in at least duplicate. Cells were stained for actin filament using Alexa Fluor™ 488-labeled phalloidin (green channel). Cells were counter stained with DAPI (Blue channel). **D**, Phalloidin Mean fluorescence intensity after EG-011 treatment (500 nM and 5 μM) at 3 time points (4, 8 and 24 h) compared to DMSO in sensitive cell line (VL51). Kruskal-Wallis test followed by Dunn’s multiple comparison was performed. **E**, Number of high fluorescence intensity spots in a sensitive lymphoma cell line after EG-011 treatment (500 nM and 5 μM) at 3 time points (4, 8 and 24h) compared to DMSO. Cells are stained with phalloidin for filamentous actin (F-actin) visualization. Ratio pared T-test was performed. **F**, Baseline levels of active WASp in sensitive and resistant cell lines. Cells are stained with antibody recognizing the active form of WASp. **G**, Increased levels of active WASp after 8 h of EG-011 treatment (500 nM and 5 μM). All experiments are performed in at least duplicate. Kruskal-Wallis test followed by Dunn’s multiple comparison was performed. Ns = nonsignificant; * p>0.05; **p<0.01; ***p<0.001; ****p<0.0001

Since the pattern of expression of WASp, expressed only in hematopoietic cells ^54,55^, was compatible with the observed activity exclusively in hematological cancers, we performed experiments to confirm WASp as target of EG-011. One of the main functions of WASp is the regulation of the actin filament nucleation activity of Arp2/3 complex ^56^. Thus, we reconstituted *in vitro* actin assembly with autoinhibited WASp, Arp2/3 complex and pyrene-labeled actin. To quantify the acceleration of actin polymerization, we used the time at which 50% of the actin is polymerized, *t_50_* (Figure 5B, right). EG-011 addition accelerates actin polymerization in presence of an autoinhibited WASp construct (Figure 5B) but had no effect on actin polymerization in the absence of autoinhibited WASp or Arp2/3 complex (Supplementary Figure 8A, B). This is consistent with a direct effect mediated by WASP (Supplementary Figure 8A, B). Moreover, EG-011 had only a modest impact on polymerization when a constitutively active WASp was employed (Supplementary Figure 8C), strongly indicating that EG-011 functions through activation of WASp.

If EG-011 activates WASp with respect to actin polymerization, then we would predict that sensitive cells might show increased actin polymerization upon EG-011 treatment. We examined cellular actin filament distribution using imaging. Cell lines were stained with Alexa Fluor 488 phalloidin following treatment with EG-011 or DMSO vehicle for 4, 8 and 24 h (500 nM and 5 μM). A significantly increase in actin polymerization was seen in EG-011 sensitive (VL51) and not in resistant (Z-138) cell lines at 4, 8 and 24 h. An increase in actin polymerization was seen as a general increase in fluorescence but also as an increase in the numbers of high intensity filamentous actin spots (Figure 5 C-E; Supplementary Figure 9). No differences were seen in the baseline levels of F-actin between sensitive and resistant cell lines

We hypothesized that baseline levels of WASp autoinhibited or activated forms differ between EG-011 sensitive and resistant cell lines. Thus, we stained sensitive (Mino, VL51) and resistant cells (MAVER1, Z138, U2932) with a specific antibody recognizing a WASp epitope available only in its activated form ^57^. We did not observe any differences in baseline level of the activated form of WASp (Figure 5F).

After exposure of a sensitive cell line (VL51) to DMSO or EG-011 for 8 h (500 nM and 5 μM) we stained for activated WASp. There was a significant increase in active WASp after EG-011 treatment, consistent with the changes in actin filament intensity stemming directly from WASp activation (Figure 5G). Thus, while the baseline degree of WASp activation does not explain the difference between sensitive and resistant cells, EG-011 does promote WASp activation.

### EG-011 *in vitro* synergizes with anti-lymphoma agents

EG-011-containing drug combinations were tested in EG-011 sensitive DLBCL (OCI-LY-1, OCI-LY-8, TMD8) and MCL (REC1, MINO) cell lines. Combination partners included the anti-CD20 monoclonal antibody rituximab, the chemotherapeutic agent bendamustine, the BCL2 inhibitor venetoclax, the BTK inhibitor ibrutinib and the immunomodulator lenalidomide, all FDA approved agents (Supplementary Figure 9). Ibrutinib and lenalidomide were tested in MCL and ABC-DLBCL (TMD8) were they showed clinical response. The synergism was assessed with Chou-Talalay combination index and as potency and efficacy according to the MuSyC algorithm ^34,35^. Based on Chou-Talalay index EG-011 showed synergism (median CI<0.9) with all the tested compounds in all the cell lines tested. Based on potency and efficacy parameters all the combinations tested showed an overall additivity or even synergism in efficacy and at lesser extent in potency, as highlighted by the majority of the points being present in the upper right part of the squares (Supplementary Figure 9 B-C). EG-011 in combination with rituximab and ibrutinib showed an effect that was in the majority of the cases beneficial in terms of efficacy while the increase in potency was given only by addition of rituximab and ibrutinib to EG-011 and not vice versa.

## Discussion

Here, we report the discovery of a novel first-in-class small molecule, EG-011, with anti-tumor activity in lymphoma, leukemia and MM models. The compound demonstrated cytotoxic activity in one third of 62 lymphoma cell lines. Cell lines belonging to the GCB-DLBCL, MZL and MCL were particularly sensitive to the treatment. The *in vitro* anti-tumor activity was confirmed in an *in vivo* experiment with a MCL xenograft, with no evidence of toxicity.

Using different approaches, we demonstrated that EG-011 targets WASp. WASp was initially identified in an unbiased screen by a change in thermal stability upon EG-011 addition. EG-011 activated WASp in reconstituted Arp2/3 complex dependent actin polymerization assays. WASp activation by EG-011 was confirmed in cells using both conformational specific antibodies and increased actin polymerization in sensitive cells. As previously mentioned, WASp is predominantly found in an autoinhibited closed conformation and gets activated by competitive binding of the small GTPase Cdc42 ^30^. The active conformation is more open and unstable, in line with the decreased stability of WASp after EG-011 treatment. Importantly, WASp expression is exclusive to blood cells, explaining EG-011’s lack of activity in a panel of cell lines derived from the most common solid tumors.

Although apparently conflicting, the anti-tumor activities described here with an activator of WASp and in published papers with a WASp inhibitor, are in line with the observed increased apoptosis and impairments of lymphocytes in individuals carrying or LOF or GOF WASp mutations ^12,13,16^. Due to the fundamental role of WASp in the regulation of the cytoskeleton and cell division in lymphocytes, cell death of blood cancer cells appears pharmacologically achievable with different classes of WASp-targeting agents. In addition to the observed direct antitumor activity, EG-011 could also have a role in the tumor microenvironment increasing T-cell capacity to kill tumor cells^58^.

Transcriptome profiling of lymphoma cells exposed to EG-011 strengthened WASp as the drug target. There was downregulation of MYC targets and of transcripts involved in mitotic spindle assembly and upregulation of genes involved in actin filament organization. Most importantly, the drugs with the most similar gene expression signatures to EG-011 were taxanes and HDAC inhibitors. Indeed, both microtubules-stabilization induced by taxanes and actin acetylation by HDAC inhibitors can lead to actin polymerization and cytoskeleton reorganization, followed by cell death ^59,60^, as seen when exposing cells to EG-011.

EG-011 showed synergism or additivity with rituximab, bendamustine, venetoclax, ibrutinib and lenalidomide, indicating its potential use also in combination with other drugs. Finally, sensitivity to EG-011 was maintained in cells with acquired resistance to other agents. EG-011 was similarly active in splenic MZL cells before and after developing resistance to PI3K and BTK inhibitors. In MM, EG-011 was more active in the cells that had become resistant to proteasome inhibitors than in their parental counterparts.

In conclusion, EG-011 is a first-in-class activator of the WASp auto-inhibited conformation displaying anti-tumor activity in lymphoma, leukemia and MM models. The data provide the rationale to further explore this mechanism of action in hematological malignancies

## Acknowledgments

We thank Jan Burkhardt and Ed Williamson (University of Pennsylvania) for the gift of the conformationally specific WASp antibody, Michael K. Rosen and Lynda Doolittle (University of Texas, Southwestern Medical Center) for the WASp expression constructs, Silvia Jenni, Yi-Chien Tsai (Children’s Hospital Zurich, Zurich, Switzerland) for their technical assistance.

## Author’s contributions

F.S., G.S., L.B., A.J.A., M.G., A.M.C.D.A., M.R.T., C.T., L.C., G.Go., R.B., M.R., S.H., K.R., F.M., G.Gu., G.V., C.D., B.B., S.B.P, N.P., E.G., P.V. performed experiments. F.S., E.G., F.B., developted the methodology. F.S., G.L., A.M.C.D.A., S.C., M.R., S.B.P, E.G., analyzed data. F.S., G.S., M.R.T., G.Gu., S.C., S.B.P, F.B., E.G., wrote and/or edited the paper. A.L., G.C., R.R., A.C., S.A., provide computational chemistry advice. F.P., E.Z., A.S., F.T., F.C., provide scientific advice. F.B., E.G., contributed to the conception or design of the study.

