## Supplementary materials and figures for "A first-in-class Wiskott-Aldrich syndrome protein (WASp) activator with anti-tumor activity in hematological cancers"

**Supplementary materials and methods**

Solid tumor and leukemia cell lines

Cell lines derived from solid tumors (Supplementary Figure 3) were obtained from the American Type Culture Collection (ATCC; Rockville, MD, USA). Cells were cultured in RPMI 1640 with L-glutamine medium (Gibco, ThermoFisher Scientific Inc., Waltham, MA, USA) supplemented with 10% FBS (#26140079, Gibco FBS qualified USA origin, Life Technologies, Saint Aubin, France), 2 mM glutamine (PAA Laboratories, Velizy-Villacoublay, France), 100 units/ml penicillin and 100 µg/ml streptomycin (PAA Laboratories). MCF7 cells were cultured in MEM Eagle (ATCC; Rockville, MD, USA) supplemented with 0.01mg/ml human recombinant insulin (Sigma-Aldrich, France), 10% FBS (#26140079, Gibco FBS qualified USA origin, Life Technologies, Saint Aubin, France), 2 mM glutamine (PAA Laboratories, Velizy-Villacoublay, France), 100 units/ml penicillin and 100 µg/ml streptomycin (PAA Laboratories). Cells lines were cultured at 37°C in a humidified 5% $CO_2$ atmosphere. Cell lines with a passage number in the range between 5 and 40 were maintained in culture for a maximum of 6 weeks.

Seven T-cell acute lymphoblastic leukemia (ALL), two B-cell ALL, one promyelocytic leukemia and one medulloblastoma cell lines were cultured in RPMI-1640 medium (Gibco, US). Medulloblastoma cell line DAOY were grown in MEMalpha and the colon cancer cell line HT29 in DMEM. Cell culture media were supplemented with fetal bovine serum (10%) (Gibco, US), Penicillin-Streptomycin (10,000 U/mL) (1%) (Gibco, US) and L-glutamine (1%) (Gibco, US). All cell lines were purchase from ATCC and all the experiment were performed within 1 month from being thawed and periodically tested for Mycoplasma negativity.

MTT proliferation assay, Cell cycle and Apoptosis assay

Lymphoma and solid tumor cell lines were exposed to a large range of concentrations of EG-011 as single agent for 72h, followed by MTT and after 4h SDS to stop the reaction. The day after, plates were read using a Cytation 3 multimode plate reader (Biotek) and $IC_{50}$s were calculated. $IC_{50}$ is defined as the drug concentration giving 50% of proliferating cells compared to vehicle treated cells. Lymphoma cell lines with $IC_{50}$ below 1µM were considered sensitive to the treatment. Area under the curve (AUC) was calculated with GraphPad.

Cell viability of twelve acute lymphoblastic leukemia (ALL) primary patient cells from different high-risk subgroups (VNN2+, E2A-HLF, refractory T and IKZF plus) co-culture with marrow–derived MSCs were assayed after 72h of incubation with EG-011 and controls.

Cell cycle and Apoptosis assay

For cell cycle evaluation, cell lines were fixed in cold 70% ethanol and stained with propidium iodide (PI) (Sigma Aldrich, US). Cells were analyzed by FACS and percentages of cells in G1, S and G2/M phases of the cell cycle were determined using the FlowJo software (TreeStar Inc., Ashland, USA).

Apoptosis was evaluated through Annexin V assay (AnnexinV-FITC apop kit, Invitrogen, US). Briefly, cells were harvested, washed with PBS and stained with Annexin V (AV) and subsequently with propidium iodide (PI). Cells AV positive/PI negative or AV positive/ PI positive were considered as apoptotic cells.

Patient-derived xenografts

Drug responses in ALL patient-derived xenografts (PDX) were analyzed as previously described [1]. Primary PDX were cultured on mesenchymal stroma cells, and drugs were added in serial concentrations. Cell viabilities were assessed by microscopy on an ImageXpress Micro microscope (Molecular Devices) after staining with CyQuant (Life Technologies, Thermo Fisher Scientific). Data were normalized against DMSO-treated controls, using a 4-parameter log logistic function (R package drc [2]).

Xenografts experiments

NOD-Scid (NOD.CB17-*Prkdcscid*/NCrHsd) mice were purchased from Harlan Laboratory (five-six weeks of age, approximately 20 g body weight). Mice maintenance and animal experiments were performed under institutional guidelines established for the animal facility and with study protocols approved by the local Cantonal Veterinary Authority (No. TI-20-2015). Mice were engrafted with the MCL REC1 cell line ($15x10^6$ cells in 100 µL of PBS). Starting with tumors of 180 $mm^3$ volume as average, mice underwent two weeks of treatment with: EG-011 (200 mg/kg, IP, 5 days per week, no.= 9 mice), control (vehicle, ip, no.= 8 mice). EG-011 was dissolved in 25% hydroxypropyl beta cyclodextrin (HP-ß-CD) in water and adjusted at pH 6.0 using NaOH (0.1 N) and HCl (0.1 N) for *in vivo* experiments. Application volume was 4,5 mL/kg (100 mg in 4.5 mL), and mice received 200 microliters (corresponding to 200 mg/kg) though intra peritoneal i.p. injection, once per day and five days per week. Tumor size was measured two times per week using a digital caliper [tumor volume ($mm^3$) = TV=D×d2/2]. Differences in tumor volumes and weights were calculated using the Mann-Whitney test (Prism 9, Version 9.3.1). The p-value for significance was $< 0.05$. The Body Condition Scoring was used to assess mice health status [3].

Thermal proteome profiling (TPP)

Experiments were performed as described in Franken et al. [4] with some modifications. Cells were resuspended in ice-cold PBS. Following resuspension, cells were homogenized in vertical tip sonication at 50% and intensity at 25%, with manual cycles of 10”/5” up to a total sonication time of 3 min, before ultracentrifugation at 100,000 g for 60 min at 4 °C to collect the soluble proteome that will be analyzed by TPP. Protein concentration was determined by BCA assay [5].The soluble proteome at 1 mg/ml was incubated with the studied drug, EG-011 at 10 µM, corresponding to the recently obtained $IC_{50}$ for this compound. The control condition following the same incubation condition with DMSO that was the vehicle solution used to solubilize the drug, The samples were incubated for 10 min at 25 ºC. For the studied EG-011, incubation was performed at the compound $IC_{50}$, and for the control, in the presence of the compound vehicle (DMSO). Seven aliquots of 100 µg of protein were individually heated for 3 min at different temperatures: 37 °C, 42 °C, 47 °C, 52 °C, 57 °C, 62 °C and 67 °C, followed by 3 min at room temperature. Subsequently, the samples were centrifugated at 100,000 g for 20 min at 4 °C. The supernatants were analyzed by label-free liquid chromatography-tandem mass spectrometry (nLC-MS/MS). In accordance with the TPP method [4], two biological replicates for the thermal shift assay were performed.

Filter Aided Sample Preparation (FASP)

Protein samples were prepared according to published methods [6]. First, the protein samples corresponding to the supernatants after centrifugation were prepared with SDT buffer (2% SDS, 100 mM Tris-HCl, pH 7.6 and 100 mM DTT), according to Wiśniewski et al.[6]. To perform FASP, the samples were diluted with 200 µl of 8 M urea in 0.1 M Tris/HCl, pH 8.5 (UA) in 30 kDa microcon centrifugal filter units. The filter units were centrifuged at 14,000 g for 15 min at 20 °C. The concentrated samples were diluted with 200 µl of UA and centrifuged at 14,000 g for 15 min at 20 °C. After discharging the flow-through 100 µl of 0.05 M iodoacetamide was added to the filter units, mixed for 1 min at 600 rpm on a thermo-mixer, and incubated static for 20 min in dark. The solution was drained by spinning the filter units at 14,000 g for 10 min. The filter units were washed three times with 100 µl buffer UA and centrifuged at 14,000 g for 15 min. The filter units were washed three times with 100 µl of 50 mM ammonium bicarbonate. Endopeptidase trypsin solution in the ratio 1:100 was prepared with 50 mM ammonium bicarbonate, dispensed, and mixed at 600 rpm in the thermomixer for 1 min. These units were then incubated in a wet chamber at 37 °C for about 16 h to achieve effective trypsinization. After 16 h of incubation, the filter units were transferred into new collection tubes. To recover the digested peptides, the tubes were centrifuged at 14,000 g for 10 min. Peptide recovery was completed by rinsing the filters with 50 µl of 0.5 M NaCl and collected by centrifugation. The samples were acidified with 10% formic acid (FA) to achieve pH between 3 and 2. The desalting process was performed by reverse phase chromatography in C18 top tips using acetonitrile (ACN; 60% v/v) with FA (0.1% v/v) for elution, and vacuum dried to be stored at −80 °C till further analysis.

Nano LC-MS/MS Analysis

The desalted peptides were reconstituted with 0.1% FA in ultra-pure milli-Q water and the concentration was measured using a Nanodrop (Thermo Scientific). Peptides were analyzed in a QExactive quadrupole-orbitrap mass spectrometer (Thermo Scientific). Samples were separated using an EASY nLC 1200 system (Thermo Scientific) and tryptic peptides were injected into a pre-column (Acclaim PepMap 100 Å, 75 um × 2 cm) and peptide separation was performed using an EASY-Spray C18 reversed-phase nano LC column (PepMap RSLC C18, 2 um, 100 Å, 75 um × 25 cm). A linear gradient of 6 to 40% buffer B (0.1% FA in ACN) against buffer A (0.1% FA in water) during 78 min and 100% buffer B against buffer A till 100 min, was carried out with a constant flow rate of 300 nl/min. Full scan MS spectra were recorded in the positive mode electrospray ionization with an ion spray voltage power frequency (pf) of 1.9 kV (kV), a radio frequency lens voltage of 60 and a capillary temperature of 275 °C, at a resolution of 30,000 and top 15 intense ions were selected for MS/MS under an isolation width of 1.2 m/z units. The MS/MS scans with higher energy collision dissociation fragmentation at normalized collision energy of 27% to fragment the ions in the collision induced dissociation mode.

Peptide and protein identification and quantification

Proteome Discoverer (v2.1, Thermo Fischer Scientific) was used for protein identification and quantification. The MS/MS spectra (.raw files) were searched by Sequest HT against the Homo sapiens UniProt database (UP000005640; 79,052 entries). A maximum of 2 tryptic cleavages were allowed, the precursor and fragment mass tolerance were 10 ppm and 0.6 Da, respectively. Peptides with a false discovery rate (FDR) of less than 0.01 and validation based on q-value were used as identified. The minimum peptide length considered was 6 and the FDR was set to 0.1. Proteins were quantified using the average of top three peptide MS1-areas, yielding raw protein abundances. Common contaminants like human keratin and bovine trypsin were also included in the database during the searches for minimizing false identifications.

Analysis of TPP experiments

Melting curves were calculated using a sigmoidal fitting approach with the R package TPP, as described in[4], with modifications. The fold changes were changed to correspond to the 7 temperatures, and the filter criteria for normalization were adjusted to this number of temperatures. The melting curves were fitted after normalization following the equation described in [7], computed in R:

$$f(T) = \frac{1 - plateau}{1 + e^{-\left(\frac{a}{T} - b\right)}} + plateau$$

where $T$ is the temperature, and a, b and "plateau" are constants. The value of f($T$) at the lowest temperature $T_{min}$ was fixed at 1. The melting point of a protein is defined as the temperature Tm at which half of the amount of the protein has been denatured. The quality criteria for filtering the sigmoidal melting curves were: (i) fitted curves for both vehicle- and compound-treated conditions had an $R^2$ of >0.8; (ii) the vehicle curve had a plateau of <0.3; (iii) the melting point differences under both the control and the treatment conditions were greater than the melting point difference between the two controls; and (iv) in each biological replicate, the steepest slope of the protein melting curve in the paired set of vehicle- and compound-treated conditions was below −0.06. The NPARC of the R package was used to detect significant changes in the temperature-dependent melting behavior of each protein due to changes in experimental conditions[4]. The significance threshold was set at $p < 0.05$.

Immunofluorescence

After treatment, cells were allowed to attach to poly-L-lysine coated slides and then fixed for 20 min with PFA 4% at room temperature (RT). Cells were permeabilized with PBS + 0.1% Triton X-100 10min at RT. To avoid unspecific staining samples were blocked for 1 hour with PBS + 5% BSA at RT before staining. To stain F-actin we used Alexafluor-488-phalloidin (Fluorescein Phalloidin, Invitrogen, US). Samples were incubated 45 min at RT. Slides were counterstained after 3 washes of PBS with 0.3 µg/mL 4,6-diamidino-2-phenylindole (DAPI) (Sigma-Aldrich). To stain for activated WASp, cells were treated the same but immunostained following fixing, permeabilizing and blocking described above. Antibodies were diluted in PBS + 5% BSA. The primary anti-'active WASp' antibody was used at 1:100 dilution. Samples were incubated overnight at 4°C. Then, samples were washed with PBS + 1% BSA and stained with a 1:1000 dilution of AlexaFluor-488 labeled goat anti-mouse secondary antibody (ab150113, Abcam, UK). Images for fluorescence intensity analysis were acquired on widefield Nikon Eclipse E800 microscope, with a 20x objective (NA 0.75). GFP and DAPI filter were used, excitation with Lumencor sola LED light and image acquisition with NIKON DS FI3 camera. Two images of more than 100 cells each were analyzed for each biological replicate. The representative images shown were acquired using a confocal microscope Leica tcs sp5. Objective HCX PL APO lambda blue, 63X/NA 1.4, oil immersion objective was used. Pixel size 80.1 nm. Excitation was performed with 405 nm diode laser and argon laser (488 nm), collection in ranges 380-493 nm and 493-655 nm respectively. Analysis was performed with ImageJ software. Briefly, images were converted in grayscale. ROI were selected using a threshold of pixel intensity of 5 and 15 for mean fluorescence intensity and high spots fluorescence intensity respectively. Roiling ball background subtraction was applied.

### Supplementary Results

#### Chemical synthesis

1-[(4-phenoxyphenyl) methyl]-3-(1-prop-2-enoyl-3-piperidyl) pyrido [3, 2-d] pyrimidine-2, 4-dione, namely EG-011 was chemical synthesized by Chimete by following a scheme reported in the Figure 1C.

Step 1: synthesis of methyl 3-[(1-tert-butoxycarbonyl-3-piperidyl)carbamoylamino] pyridine-2-carboxylate

To a solution of 3-isocyanatepyridine-2-carboxy1ate (prepared as described in Synthetic Communications, 2003, 33(24), 4259-4268; 1.20 g, 6.736 mmol) in dry dichloromethane (8 mL), a solution of racemic tert-butyl 3-aminopiperidine-1- carboxylate (1.349 g, 6.736 mmol) in DCM (4 mL) was added drop-wise and the resulting mixture was stirred at room temperature overnight. After evaporation of the solvent at reduced pressure, the residue was purified by flash chromatography on Biotage KP-Sil SNAP cartridge (DCM : EtOAc = 80 : 20 to 100% EtOAc). A further purification by flash chromatography on Bitotage NH SNAP cartridge (hexane : EtOAc = 80 : 20 to 40 : 60) was required to afford title compound (1.4 g). MS/ESI+ 379.1 [MH]+, Rt = 13.6 min

Step 2: synthesis of tert-butyl 3-(2,4-dioxo-1H-pyrido[3,2-d]pyrimidin-3-yl)piperidine- 1-carboxylate.

To a solution of methyl 3-[(1-tert-butoxycarbonyl-3 piperidyl)carbamoylamino]pyridine-2-carboxylate (1.3 g) in MeOH (15 mL), 20% sodium ethoxide solution in EtOH (1.48 mL, 3.8 mmol) was added and the resulting mixture was heated to reflux for 8 h, then stirred at room temperature overnight. The mixture was evaporated to dryness together with a smaller batch (obtained by reacting 0.100 g of methyl 3-[(1-tert-butoxycarbonyl-3-piperidyl)carbamoylamino]pyridine-2-carboxylate under the same conditions) and the residue was dissolved in water and acidified with acetic acid (pH ≈ 4-5). The precipitate was collected by filtration and washed several times with plenty of water.

The solid was dried under vacuum at 50°C affording title compound as a beige solid (1.1 g, 3.175 mmol, 47% yield over 2 steps). MS/ESI+ 347.0 [MH]+, Rt = 11.8 min

Step 3: synthesis of tert-butyl 3-[2,4-dioxo-1-[(4-phenoxyphenyl)methyl]pyrido[3,2- d]pyrimidin-3-yl]piperidine-1-carboxylate.

To a suspension of tert-butyl 3-(2,4-dioxo-1H-pyrido[3,2-d]pyrimidin-3-yl)piperidine-1-carboxylate (1.00 g, 2.89 mmol) in dry DMF (8 mL), $K_2CO_3$ (0.798 g, 5.78 mmol) was added followed by a solution of 1-(bromomethyl)-4-phenoxy-benzene (0.911 g,3.46 mmol) in DMF (1 mL) and the resulting mixture was stirred at room temperature for 46 h. The mixture was partitioned between EtOAc (70 mL) and water (60 mL). The organic phase was washed with brine (3 x 70 mL), dried over sodium sulfate, filtered and concentrated. The residue was purified by flash chromatography on Biotage KP-Sil SNAP cartridge (100% EtOAc) to afford title compound as a beige foam (1.380 g, 2.61 mmol, 90% yield). MS/ESI+ 529.1 [MH]+, Rt = 25.4 min

Step 4: synthesis of 1-[(4-phenoxyphenyl)methyl]-3-(3-piperidyl)pyrido[3,2-d]pyrimidine-2,4-dione

To a solution of tert-butyl 3-[2,4-dioxo-1-[(4-phenoxyphenyl)methyl]pyrido[3,2- d]pyrimidin-3-yl]piperidine-1-carboxylate (1.375 g, 2.60 mmol) in DCM (20 mL) cooled to 3°C, TFA (1.992 mL, 26.0 mmol) was added drop-wise and the resulting mixture was left to warm to r.t. and stirred for 6 h. The mixture was diluted with DCM and washed with sat. $NaHCO_3$. The organic phase was dried over sodium sulfate, filtered and concentrated to afford title compound as a beige foam (1.086 g, 2.53 mmol, 97% yield) which was used without purification. MS/ESI+ 429.0 [MH]+, Rt =8.9 min

Step 5: synthesis of 1-[(4-phenoxyphenyl)methyl]-3-(1-prop-2-enoyl-3- piperidyl)pyrido[3,2-d]pyrimidine-2,4-dione

A solution of 1-[(4-phenoxyphenyl)methyl]-3-(3-piperidyl)pyrido[3,2-d]pyrimidine- 2,4-dione (1.08 g, 2.52 mmol) and TEA (1.054 mL, 7.56 mmol) in DCM (20 mL) was cooled to 3°C and acryloyl chloride (0.246 mL, 3.02 mmol) was added. The mixture was left to warm to room temperature (RT). and stirred overnight. The mixture was diluted with DCM and washed with 5% aqueous citric acid. The aqueous phase was extracted with DCM and the combined organic layers were washed with brine, dried over sodium sulfate, filtered and concentrated. The residue was purified by flash chromatography on Biotage KP-Sil SNAP cartridge (DCM to DCM : MeOH = 95 : 5). A further purification by flash chromatography on Biotage Ultra silica SNAP cartridge (EtOAc to EtOAc:MeOH = 95 : 5) was required to afford title compound as white foam (0.640 g, 1.326 mmol, 53% yield). MS/ESI+ 483.1 [MH]+, Rt = 17.4 min; 1H NMR (300 MHz, DMSO- d6) δ ppm 8.54, (d, 1 H), 7.60 - 7.85 (m, 2 H), 7.27 - 7. 50 (m, 4 H), 7.13 (t, 1 H), 6.65- 7.05 (m, 5 H), 6.14 (d, 1 H), 5.69 (d, 1 H), 5.20 - 5.48 (m, 2 H), 4.71 - 4.88 (m, 1H), 4.40 - 4.65 (m, 1 H), 4.05 - 4.260 (m, 1 H), 3.50 - 4.05 (m, 1 H), 2.35 - 3.15 (m, 2 H), 1.77 - 1.95 (m, 2 H), 1.34 - 1.62 (m, 1 H).

### Supplementary Figures

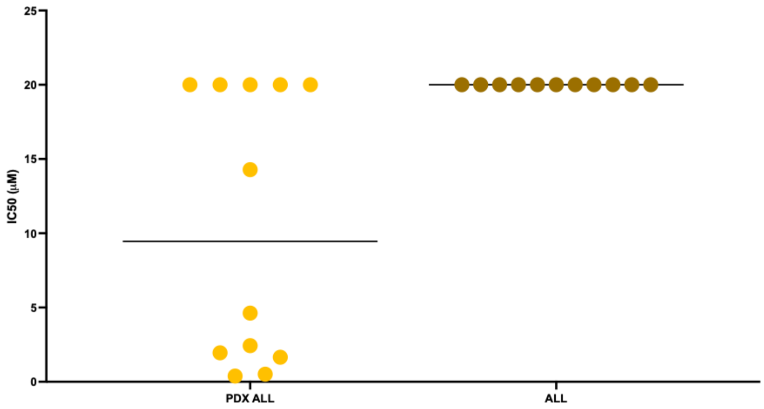


| Histology | Cell lines |
|---|---|
| T-ALL | RPMI8402 |
| T-ALL | Molt-4 |
| T-ALL | ALL-SIL |
| T-ALL | DND4;1 |
| T-ALL | TALL-1 |
| T-ALL | P12-ichikawa |
| T-ALL | CCRF-CEM |
| B-ALL | RCH-ACV |
| B-ALL | AT-2 |
| B-ALL | Nalm-6 |
| promyelocytic leukaemia | HL-60 |

**Supplementary figure 1. EG-011 activity in acute lymphoblastic leukemia (ALL) PDX and cell lines.** EG-011 in vitro activity in ALL PDX and ALL cell lines. IC50s calculated after 72h of treatment. IC50 >10 µM, arbitrary set as 20 µM. Table on the right shows ALL cell lines.

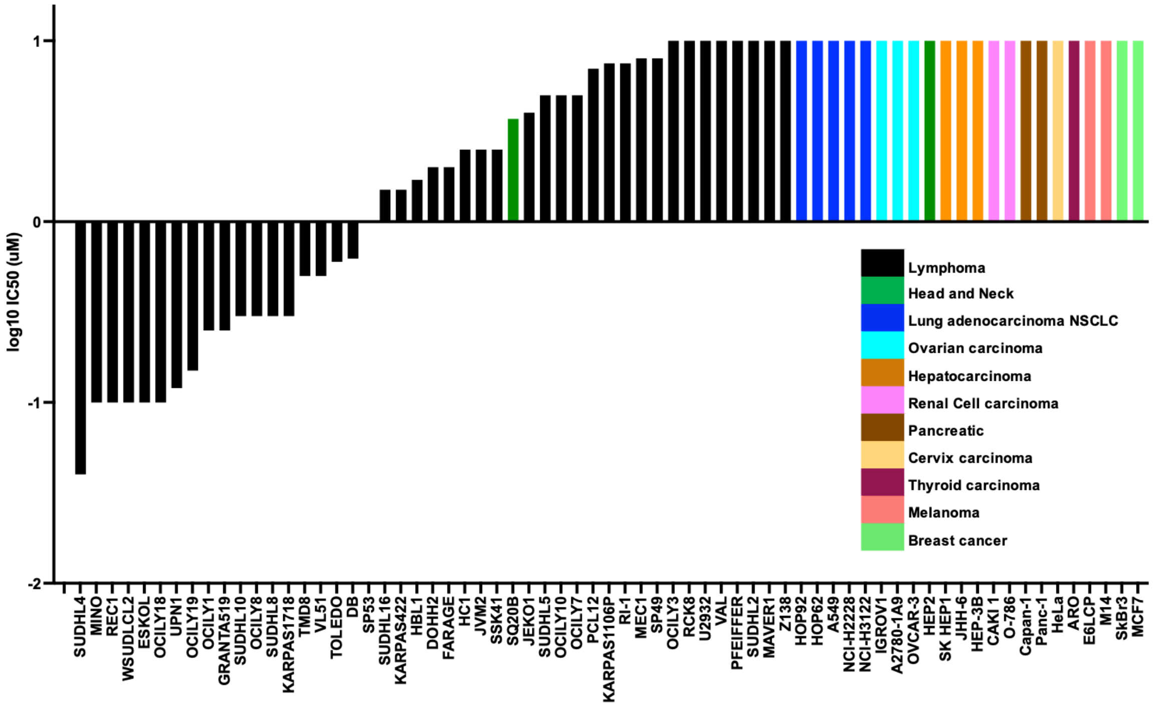


**Supplementary Figure 2. EG-011 has anti-tumor activity only in hematological cancers.** EG-011 in vitro activity in lymphoma cell lines and solid tumor cell lines. IC50s calculated after 72h of treatment. IC50 >10 μM, arbitrary set as 20 μM

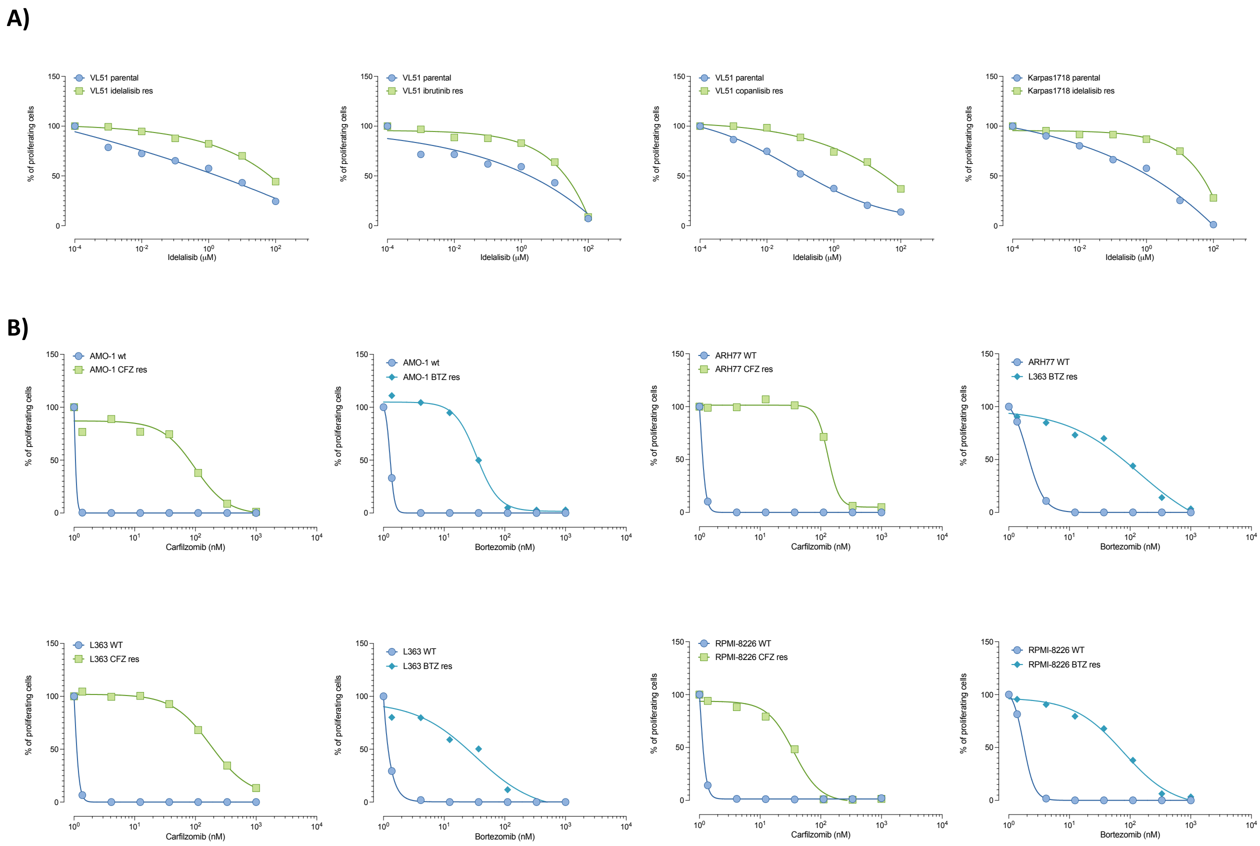


**Supplementary Figure 3. Marginal zone lymphoma (MZL) and multiple myeloma (MM) models with acquired resistance to FDA approved compounds.** A) Dose response curves after 72h treatment with idelalisib, ibrutinib or copanlisib in MZL cell lines with acquired resistance to PI3K and BTK compared to parental cells. B) Dose response curves after 72h treatment with carfilzomib (CFZ) and bortezomib (BTZ) in MM cell lines with acquired resistance to proteasome inhibitors compared to parental cells. Raw data for MZL cell lines treated with PI3K and BTK inhibitors were already published and were re-analized for the figure [8-11].

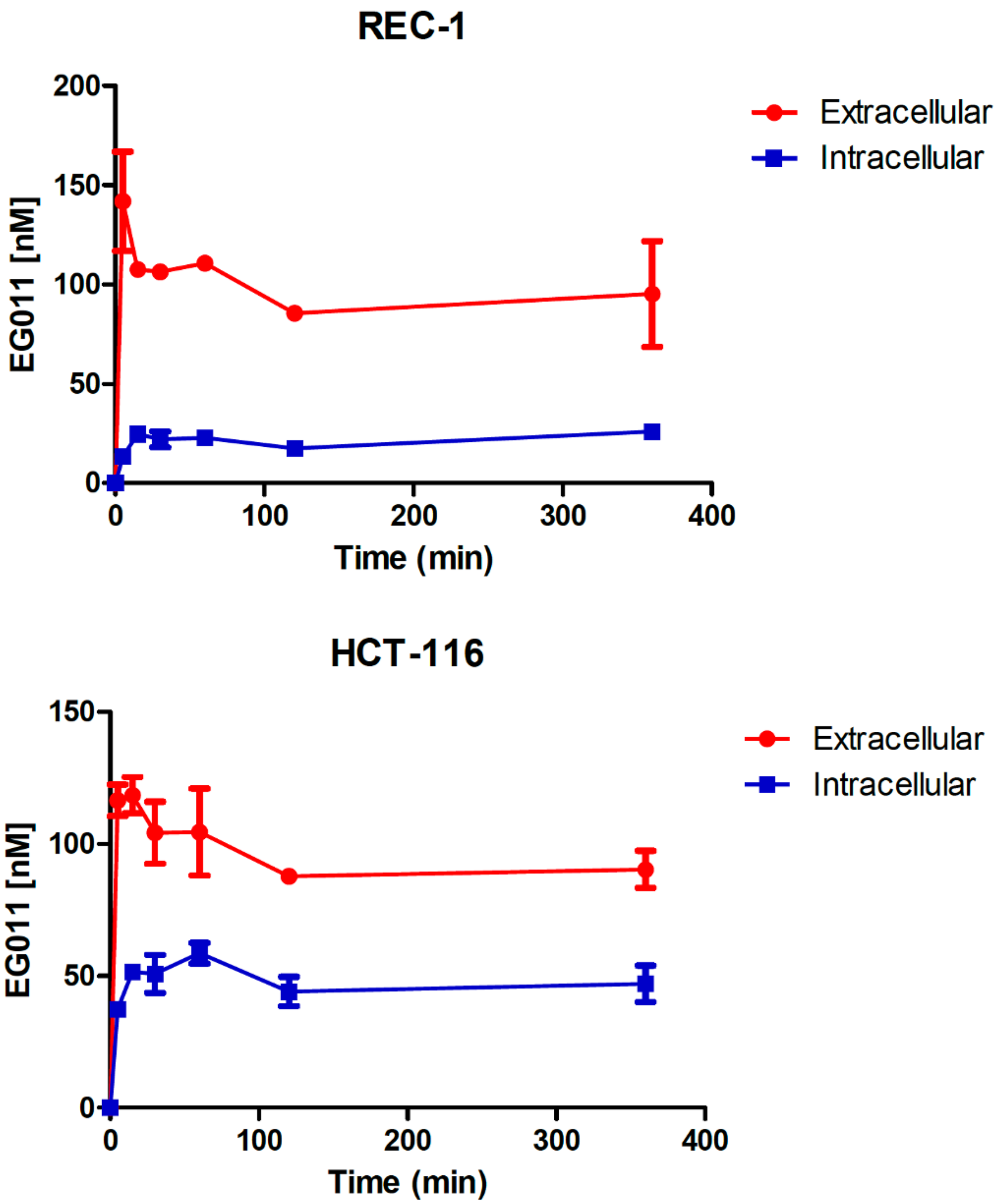


**Supplementary Figure 4. Differences in cellular uptake do not explain the lack of activity in solid tumors.** Extracellular and intracellular levels of EG-011 analyzed over time in the solid tumor resistant model HCT-116 (colorectal cancer cell line) and in the lymphoma sensitive model cell line REC1 (MCL cell line).

**L1000**

| Drug | MOA | NES |
|---|---|---|
| Nsc-632839 | ubiquitin specific protease inhibitor | 2.3529 |
| Scriptaid | HDAC inhibitor | 2.3327 |
| Entinostat | HDAC inhibitor | 2.2736 |
| Nsc-3852 | HDAC inhibitor | 2.2619 |
| Belinostat | HDAC inhibitor | 2.2492 |
| Dacinostat | HDAC inhibitor | 2.2391 |
| Cd-437 | retinoid receptor agonist | 2.2268 |
| Tw-37 | BCL inhibitor | 2.2125 |
| Bibr-1532 | telomerase inhibitor | 2.2087 |
| Bortezomib | proteasome inhibitor | 2.2029 |
| Trichostatin-A | HDAC inhibitor | 2.1987 |
| Cerulenin | fatty acid synthase inhibitor | 2.1744 |
| Tg-101348 | FLT3 inhibitor\|JAK inhibitor | 2.1508 |
| Pyroxamide | HDAC inhibitor | 2.143 |
| Panobinostat | HDAC inhibitor | 2.1384 |
| Vorinostat | HDAC inhibitor | 2.1374 |
| Doxorubicin | topoisomerase inhibitor | 2.136 |
| Ikk-2-Inhibitor-V | IKK inhibitor\|NFkB pathway inhibitor | 2.1319 |
| Gw-405833 | cannabinoid receptor agonist | 2.1267 |
| Mg-132 | proteasome inhibitor | 2.1263 |
| Menadione | mitochondrial DNA polymerase inhibitor | 2.1155 |
| Nsc-663284 | CDC inhibitor | 2.0998 |
| Serdemetan | MDM inhibitor | 2.0937 |
| Perhexiline | carnitine palmitoyltransferase inhibitor | 2.0845 |
| Ryuvidine | histone lysine methyltransferase inhibitor | 2.0783 |
| Cladribine | adenosine deaminase inhibitor | 2.0777 |
| Tyrphostin-Ag-1478 | EGFR inhibitor | 2.0583 |
| Ikk-16 | IKK inhibitor | 2.0467 |
| Tyrphostin-A9 | tyrosine kinase inhibitor | 2.042 |
| Bi-2536 | PLK inhibitor | 2.0369 |
| Sn-38 | topoisomerase inhibitor | 2.0359 |
| Elesclomol | oxidative stress inducer | 2.0358 |
| Purvalanol-A | CDK inhibitor | 2.0322 |
| Pac-1 | caspase activator | 2.029 |
| Cd-1530 | retinoid receptor agonist | 2.028 |
| Etoposide | topoisomerase inhibitor | 2.0273 |
| Quiflapon | leukotriene synthesis inhibitor | 2.0174 |
| Bx-795 | IKK inhibitor | 2.0173 |
| Calcipotriol | vitamin D receptor agonist | 1.9915 |

**GDSC**

| Drug | MOA | NES |
|---|---|---|
| Docetaxel | Microtubulin-stabilizing agent | 2.1683 |
| CGP-60474 | Cell Cycle (Interphase), Mitosis | 2.0369 |
| SL 0101-1 | ERK Signalling, Mitosis | 2.0212 |
| Bortezomib | proteasome inhibitor | 1.9013 |
| TGX221 | NA | 1.8931 |
| Epothilone B | Microtubulin-stabilizing agent | 1.8819 |
| JNK Inhibitor VIII | Stress Pathways | 1.8689 |
| BIBW2992 | ERK Signalling, PI3K/MTOR | 1.8465 |
| BMS-509744 | Adhesion, Cytoskeleton | 1.8373 |
| Gefitinib | ERK Signalling, PI3K/MTOR | 1.8287 |
| AS601245 | Stress Pathways | 1.8193 |
| RO-3306 | Mitosis | 1.8027 |
| Vinblastine | Microtubulin-destabilizing agent | -1.7079 |
| ATRA | Transcription | -1.7447 |
| MK-2206 | Apoptosis, Metabolism, PI3K/MTOR | -1.7641 |
| AZD-2281 | DNA Repair, Other | -1.7972 |
| Bosutinib | Cytoskeleton, ERK signalling, Other | -1.8322 |
| KU-55933 | DNA Repair | -1.8345 |
| AZD7762 | Cell Cycle (Interphase), DNA Repair | -1.86 |
| Salubrinal | Other | -1.8819 |
| Nilotinib | Cytoskeleton | -1.9032 |
| IPA-3 | Cytoskeleton, | -1.9898 |
| AICAR | AMPK, Metabolism | -1.9951 |
| Axitinib | Angiogenesis, ERK Signalling, PI3K/MTOR | -2.0347 |
| PD-0332991 | Cell Cycle (Interphase) | -2.0364 |
| AP-24534 | Cytoskeleton | -2.0853 |
| LAQ824 | NA | -2.0861 |
| AZD8055 | Metabolism, PI3K/MTOR | -2.1018 |
| BX-795 | Mitosis, NFkappB, PI3K/MTOR | -2.227 |
| Vorinostat | Chromatin Modification | -2.2298 |
| Methotrexate | Replication, Transcription | -2.2401 |
| ZM-447439 | Mitosis | -2.4763 |
| ABT-263 | Apoptosis | -2.5799 |

**Supplementary Figure 5. EG-011 determines transcriptome changes similar to microtubules-stabilizing agents and HDAC inhibitors**. Drug connectivity enrichment plots showing the compounds inducing similar or opposite transcriptional changes to REC1 cell line exposed to EG-011 treatment or to DMSO, as control. Transcriptional changes induced by EG-011 treatment were compared to transcriptional changes induced by other drugs present in publicly available data such as the LINCS L1000 project (L1000) and the Genomics of Drug Sensitivity in Cancer (GDSC) project.

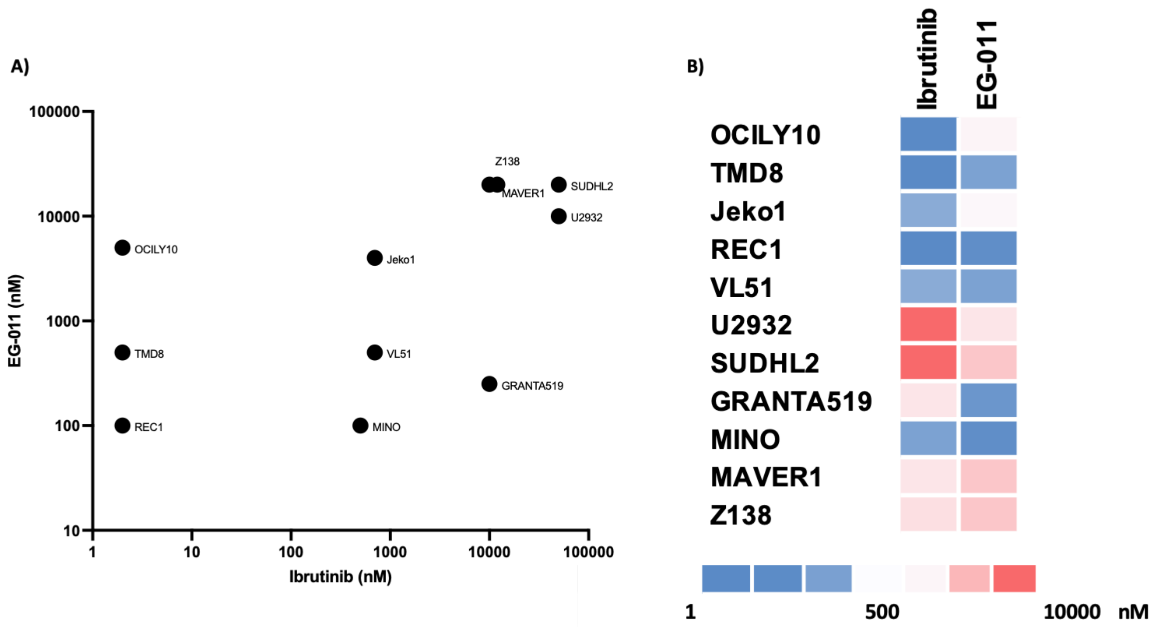


**Supplementary Figure 6. EG-011 pattern of activity differs from ibrutinib. A, I**C50s correlation in different lymphoma cell lines treated with EG-011 (y axis) and ibrutinib (X axis). **B**, Heatmap with EG-011 and Ibrutinib IC50s in lymphoma cell lines

A)

**Stabilized Protein**

| Gene name | FC Tm (Tm Treated-Tm ctr) | p-value |
|---|---|---|
| CUTC | 2.435 | 0.00030 |
| AIMP1 | 4.18 | 0.00120 |

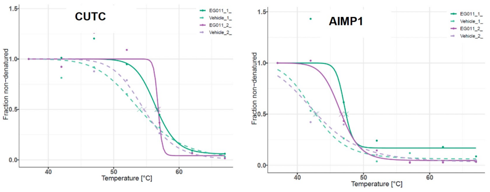


B)

**Destabilized Protein**

| Gene name | FC Tm (Tm Treated-Tm ctr) | p-value |
|---|---|---|
| WAS | -12.305 | 0.00003 |
| CAMK1D | -10.205 | 0.02700 |
| NUDCD2 | -7.125 | 0.01400 |
| ELOC | -6.35 | 0.00100 |
| C12orf10 | -6.23 | 0.02300 |
| NSUN2 | -5.43 | 0.00800 |
| ARHGAP25 | -5.255 | 0.03600 |
| SCRN1 | -4.71 | 0.00080 |
| PPIF | -4.7 | 0.01600 |
| CCT2 | -4.23 | 0.01700 |
| USP14 | -4.17 | 0.04700 |
| GSPT1 | -3.525 | 0.01000 |

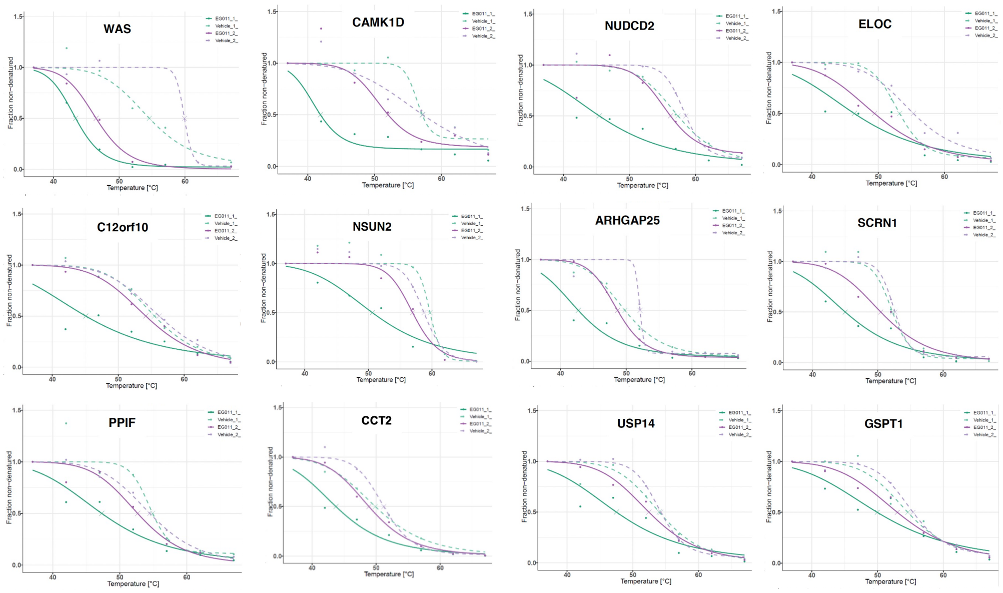


**Supplementary Figure 7. Stabilized and destabilized proteins in the presence of EG-011.** Statistically significantly stabilized **(A)** and destabilized **(B)** proteins with thermal proteomic profiling (TPP) in the presence of EG-011 or DMSO. Dashed lines, DMSO treatment; Solid lines, EG-011 treatment. Experiments performed in two replicates.

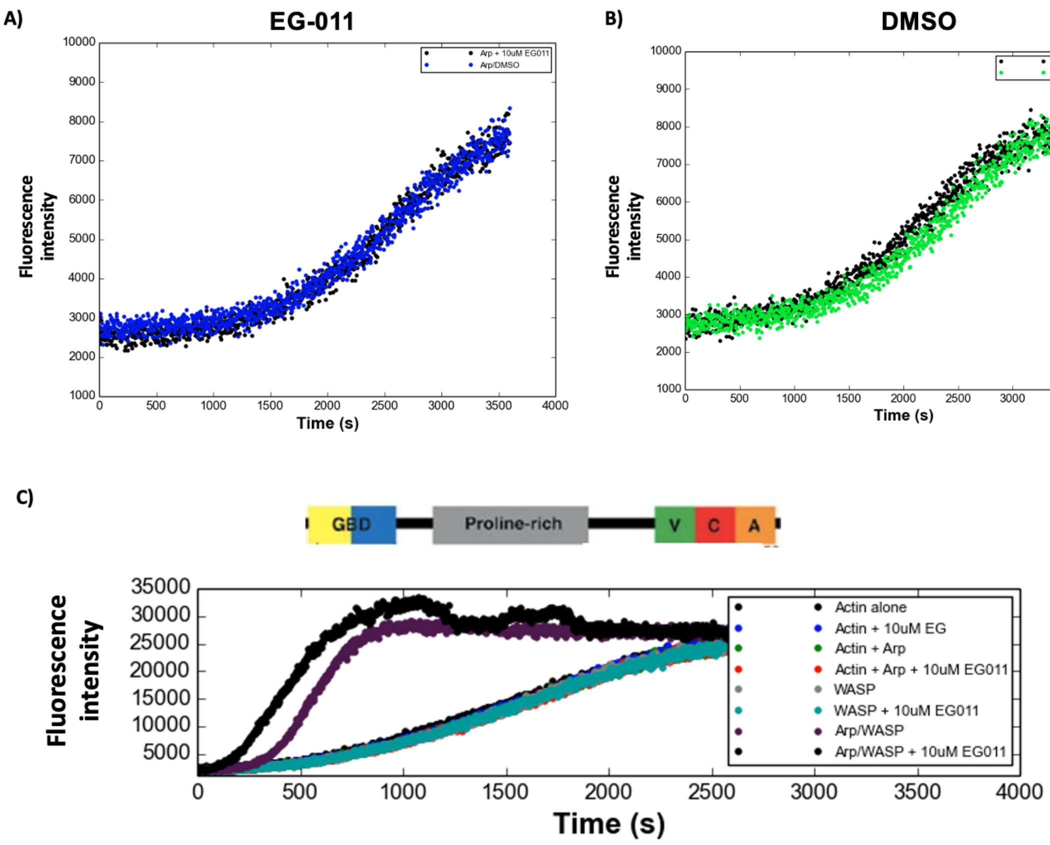


**Supplementary Figure 8. Pyrene actin polymerization controls.** Pyrene actin polymerization measured as fluorescence intensity, in the absence of WASP and treated with EG-011 (**A**) or with WASp and treated with DMSO (**B**). (**C**) Pyrene actin polymerization measured as fluorescence intensity in the absence or presence of EG-011 at 10 μM and active WASp.

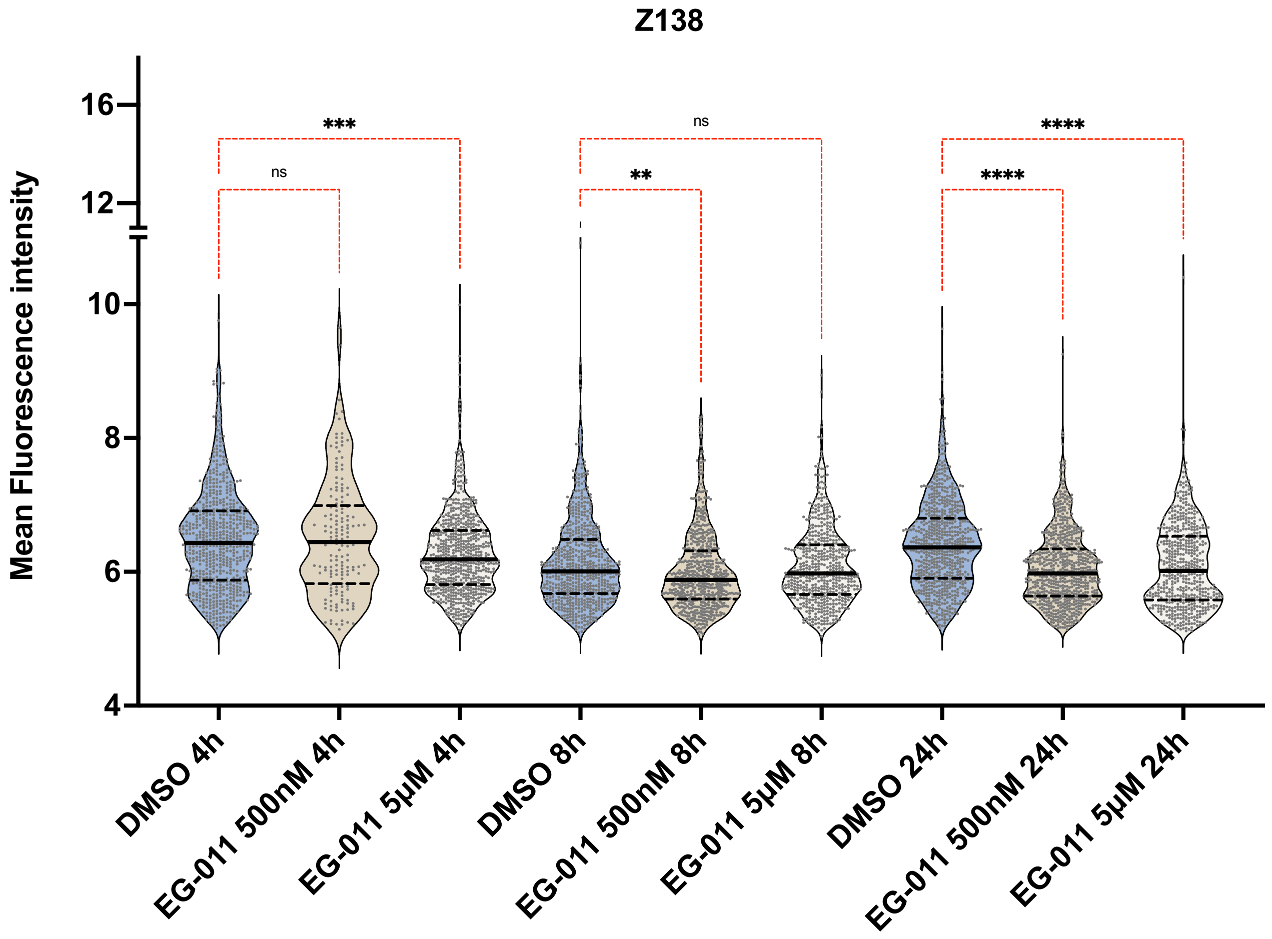


**Supplementary Figure 9. Phalloidin mean fluorescence intensity in resistant cell line.** Phalloidin mean fluorescence intensity after EG-011 treatment (500 nM and 5 μM) at three time points (4, 8 and 24h) compared to DMSO in resistant cell line Z138. Kruskal-Wallis test followed by Dunn's multiple comparison was performed. Ns = nonsignificant; **$p<0.01$; ***$p<0.001$; ****$p<0.0001$

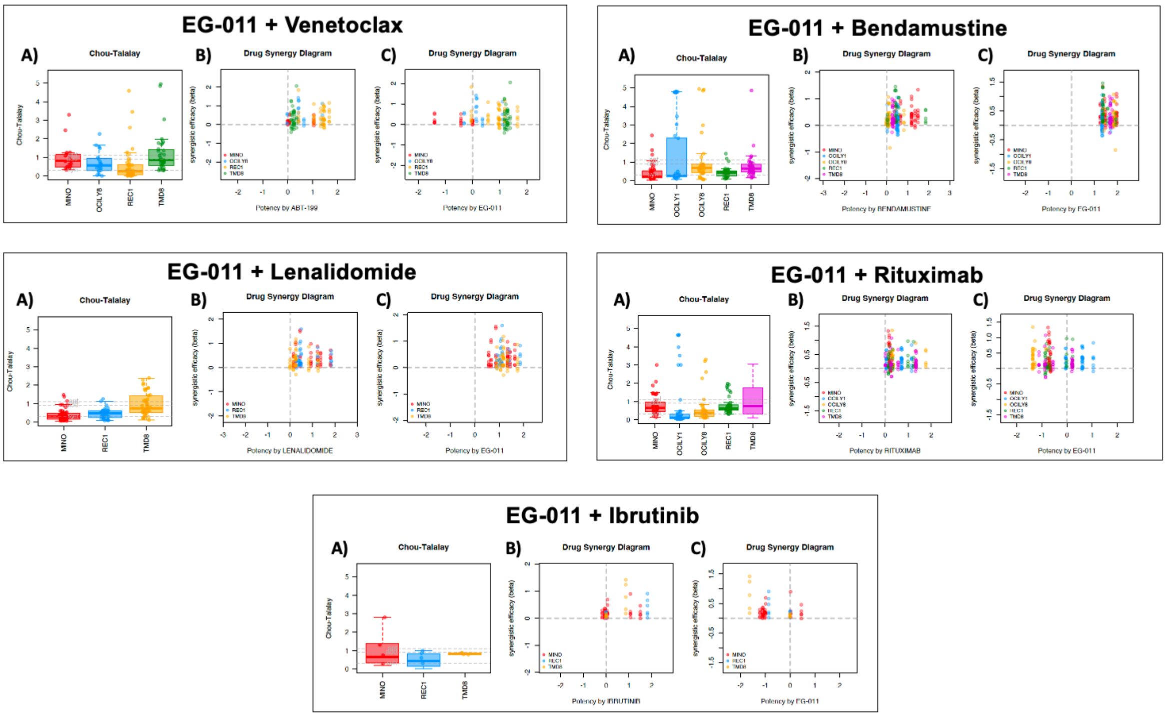


**Supplementary Figure 10. EG-011 synergizes with FDA approved compounds.** Chou-Talalay index, efficacy and potency parameters calculated for each lymphoma cell line and each combination. In each box-plot, the line in the middle of the box represents the median and the box extends from the 25th to the 75th percentile (interquartile range, IQ); the whiskers extend to the upper and lower adjacent values (i.e., 1.5 IQ). Each point represents the CI (**A**) or efficacy and potency (**B-C**) data at the different combination of concentrations in each cell line. Chou-Talalay Combination Index (CI): Synergism, CI < 0.9; additive effect, 0.9 < CI < 1.1; no benefit, CI >1.1. Efficacy: Synergism > 1; additive effect, from 0 to 1; no benefit from 0 to -1; antagonism < -1. Potency: Synergism > 0.5; additive effect, from 0 to 0.5; no benefit from 0 to 0.5; antagonism < -0.5

**Supplementary Table 1. List of lymphoma cell lines with relative histology and IC50 to EG-011**

**Supplementary Table 2. Genes and pathways modulated after EG-011 treatment.**

**Supplementary Table 3. List of compounds present in publicly available data of other drugs (L1000 and GDSC) with similar or opposite gene modulation to EG-011.**

**Supplementary Table 4. List of kinases tested as possible target of EG-011 by KINOMEscan assay PanQinase Activity Assay.** Positive hits with **%** compared to CTR less then 50%.

**Supplementary Table 5. List of proteins stabilized or destabilized by EG-011 identified with thermal proteome profiling.**
